# Mitochondrial function is essential for humoral immunity by controlling flux of the TCA cycle, phosphatidic acid and mTOR activity in B cells

**DOI:** 10.1101/2021.01.14.426649

**Authors:** Sophia Urbanczyk, Olivier R. Baris, Jörg Hofmann, Florian Golombek, Kathrin Castiglione, Xianyi Meng, Aline Bozec, Dimitrios Mougiakakos, Sebastian R. Schulz, Wolfgang Schuh, Ursula Schlötzer-Schrehardt, Tobit D. Steinmetz, Susanne Brodesser, Rudolf J. Wiesner, Dirk Mielenz

## Abstract

The function of mitochondrial respiration during B cell fate decisions and differentiation remains equivocal. This study reveals that selection for mitochondrial fitness occurs during B cell activation and is essential for subsequent plasma cell differentiation. By expressing a mutated mitochondrial helicase in transitional B cells, we depleted mitochondrial DNA during B cell maturation, resulting in reduced oxidative phosphorylation. Although no changes in follicular B cell development were evident, germinal centers, class switch recombination to IgG, plasma cell generation and humoral immunity were diminished. Defective oxidative phosphorylation led to aberrant flux of the tricarboxylic acid cycle and lowered the amount of saturated phosphatidic acid. Consequently, MTOR activity and BLIMP-1 induction were curtailed whereas HIF1α, glycolysis and AMPK activity were amplified. Exogenous phosphatidic acid increased mTOR activity in activated B cells. Hence, mitochondrial function is required and selected for in activated B cells for the successful generation of functional plasma cells.

## Introduction

Humoral immunity depends on the development of antibody (Ab) secreting long-lived plasma cells. There are T cell independently (TI) generated, IgM secreting short-lived plasma cells derived from marginal zone (MZ) or B1 B cells, T cell dependently (TD) activated and class switched plasma cells secreting mainly IgA in gut associated tissues or long lived plasma cells secreting IgA, IgM or IgG in the bone marrow (Schuh et al., 2020). All plasma cells have undergone profound anabolic and morphologic changes, such as cell growth and expansion of the endoplasmic reticulum during several cell divisions (Nutt et al., 2011). Several models explain the incremental up-regulation of plasma cell specific transcription factors of the B cell lineage, such as BLIMP1 or XBP1 (Hawkins et al., 2013). BLIMP1 is required for the fully secretory phenotype of plasma cells (Kallies et al., 2007) which appears to involve oxidative phosphorylation (OxPhos) (Price et al., 2018). On the other hand, mature plasma cells depend on glucose uptake for Ab glycosylation and import of pyruvate into mitochondria (Lam and Bhattacharya, 2018). The metabolism of mature plasma cells is distinct from resting B cells which primarily consume fatty acids (FA) to support OxPhos in mitochondria (Caro-Maldonado et al., 2014) and depend on continuous degradation of hypoxia inducible factor 1α (HIF1α) (Xu et al., 2019). Yet, resting B cells increase OxPhos, glucose uptake and glycolysis when activated by the B cell receptor (BCR) or the innate stimulus lipopolysaccharide (LPS) already within 6h, switching to glucose and glutamine catabolism (Caro-Maldonado et al., 2014). Anti CD40/IL-4 activated B cells utilize glucose to support the pentose phosphate pathway (PPP) that is also used by many tumour cells for the rapid generation of nucleotides and other anaplerotic precursors required for cell proliferation (Waters et al., 2018). There also appears to be mitochondrial remodeling (Waters et al., 2018), with IRF4 supporting mitochondrial homeostasis in established plasma cells (Low et al., 2019). OxPhos is fueled by the Krebs/tricarboxylic acid cycle (TCA) in mitochondria (Kennedy and Lehninger, 1949). The TCA cycle utilizes progressive transformation of oxaloacetate into TCA intermediates such as citrate, α-ketoglutarate, succinate, fumarate or malate by addition or removal of C atoms. In some of these reactions, net energy is transferred to reduction equivalents, such as nicotinamide adenine dinucleotide (NAD) or flavine adenine dinucleotide (FAD), to yield NADH or FADH2 (Krebs, 1948). Complex I oxidizes NADH and while the electrons are transported alongthe electron transport chain (ETC) complexes, protons are pumped across the inner mitochondrial membrane (IMM). Complex II oxidizes FADH2 without proton transport. Complex III and IV again pump protons across the IMM while transferring electrons to O_2_, the terminal essential electron acceptor, which is reduced to H_2_O. Complex V, the ATP synthase, utilizes the chemiosmotic gradient to efficiently generate ATP. OxPhos up-regulation accompanies plasma cell generation and is maintained by IRF4 in plasma cells (Low et al., 2019; Price et al., 2018), but a causal relation between OxPhos, B cell activation and plasma cell generation has been questioned (Milasta et al., 2016). The upregulation of OxPhos genes by BLIMP1 affects nuclear genes essential for the assembly of complexes I-V of the mitochondrial respiratory chain (mtRC) (Price et al., 2018). Nonetheless, only Complex II is encoded entirely by nuclear genes, while the essential subunits of the mtRC complexes I, III, IV and V are encoded by mitochondrial DNA (mtDNA) (Gustafsson et al., 2016). In total, mtDNA codes for these 13 crucial subunits and also contains the genes for 22 tRNAs and 2 ribosomal RNAs necessary for their synthesis in the mitochondrial matrix. Expression of mtRC subunits requires continuous replication and transcription of mtDNA which needs to be unwound by the essential mitochondrial helicase PEO1/TWINKLE (TWINKLE) (Milenkovic et al., 2013; Spelbrink et al., 2001). Dominant negative mutants of TWINKLE impair mtDNA replication, thereby causing mitochondrial disease in humans due to multiple mtDNA deletions and/or depletion (Goffart et al., 2009; Sarzi et al., 2007; Spelbrink et al., 2001). Those large mtDNA deletions, even in the presence of wild-type (WT) molecules (called a heteroplasmic state), reduce the amount of newly synthesized subunits. Truncated or fused subunits further impair ETC function and ATP production (Hornig-Do et al., 2012; von Kleist-Retzow et al., 2007).

The mammalian target of Rapamycin (MTOR) complex, specifically the Rapamycin-sensitive MTORC1 complex (Iwata et al., 2017), has been proposed to integrate the metabolic fate of glucose, glutamine and lipids (Foster et al., 2014). Intriguingly, MTORC1 is required for the generation, but not maintenance, of plasma cells (Benhamron et al., 2015; Jones et al., 2016). These data suggest that there is an anabolic control of plasma cell development via MTOR (Lam and Bhattacharya, 2018). MTORC1 is activated by the AKT/Phosphoinositol-3-kinase (PI3K) pathway, by glucose, glutamine and growth factors (Iwata et al., 2017). MTORC1 is also directly activated by phosphatidic acid (PA) (Toschi et al., 2009) that is either generated through cleavage of membrane lipids via the phospholipase D (PLD) pathway or synthesized *de novo* via the lysophosphatidic acid (LPA) acyltransferase (LPAAT) (Foster et al., 2014) pathway. The LPAAT pathway integrates metabolites of glycolysis, glyeraldehyde-3-phosphate (G3P), and the TCA cycle, namely citrate, to form FA and then PA (Foster, 2013). PA is present in low amounts, acts mainly as a signaling molecule but is also a precursor in the synthesis of other membrane lipids (Athenstaedt and Daum, 1999).

We hypothesized that depletion of mtDNA in mature B cells will compromise OxPhos in activated B cells and thereby bring new insights on its function in this cell type. The approach of mtDNA depletion circumvents the use of inhibitors in mice and in cell culture. Interestingly, the murine K320E dominant negative TWINKLE (DNT) variant (Baris et al., 2015), equivalent to the K319E human mutation (Hudson et al., 2005), leads to massive mtDNA depletion when expressed in rapidly dividing cells such as keratinocytes (Weiland et al., 2018). To determine the importance of mtDNA and the mtRC in rapidly dividing B cells *in vivo,* we specifically expressed DNT in B cells to induce mtDNA depletion. We found increased replication of mtDNA during wild type B cell activation. Altered mtDNA replication in the presence of DNT hindered GC B cell development, BLIMP1 expression as well as plasma cell generation due to lowered OxPhos activity, reduced generation of PA as a consequence of the disturbed TCA cycle, and reduced MTOR activity in early plasmablasts. In addition, plasma cells expressing DNT were outcompeted. Hence, our results place OxPhos upstream of BLIMP1 and MTOR induction during plasma cell differentiation to ensure selection for mitochondrial fitness.

## Results

### Mitochondrial DNA is up-regulated in activated and marginal zone B cells

To determine the abundance of mitochondrial DNA (mtDNA) we isolated B cell subsets by flow cytometric cell sorting (Figure S1). Abundance of the *16s* rRNA gene encoded by mtDNA was quantified by qPCR and normalized to nuclear DNA (Figure 1a). MZ B cells, splenic plasma cells and BM plasma cells identified by CD138 and TACI staining (Pracht et al., 2017) exhibit higher amounts of mtDNA than follicular B cells (FO). These data suggest that there is replication of mtDNA during B cell maturation from transitional B cells to MZ B cells (Loder et al., 1999) and from resting B cells to plasma cells. GC B cells showed only a slight increase of mtDNA compared to FO B cells. GC B cells are the most rapidly dividing cells in the mammalian body (MacLennan, 1994). This implies that proliferation is accompanied by mtDNA replication in order to keep the copy number constant. To test rigorously whether proliferation and differentiation indeed increase mtDNA copy number in B cells, we quantified it in LPS-activated B cells (Figure 1b). This revealed a ∼ two-fold increase in mtDNA between d0 and d3 of activation (Figure 1b). Together, these data suggest that there is an increased requirement for proteins of the mtRC that are encoded by mtDNA (Gustafsson et al., 2016) during B cell proliferation and/or differentiation. Thus, to interfere with the function of the mtRC in activated B cells, but avoid affecting the metabolic checkpoints of early B cell development (Urbanczyk et al., 2018), we crossed mice carrying a loxP-flanked STOP cassette upstream of K320E-TWINKLE (DNT) coupled to IRES-GFP (Baris et al., 2015) with CD23CRE mice (Kwon et al., 2008) to obtain DNTxCD23CRE (short: DNT mice) (Figure 1c). We found expression of DNT, indicated by GFP expression, in B cells but not T cells (Figure 1d). To determine the onset of GFP expression we analyzed B cells in the BM (Figure S2), revealing that ∼20% of transitional B cells and ∼90% of mature, recirculating B cells express GFP. These data are consistent with the onset of CD23 expression in transitional type 2 B cells (Loder et al., 1999) and show that CRE mediated excision of the floxed STOP cassette of DNT mice does not show a complete penetrance, enabling competition of STOP cassette-deleted vs. -non-deleted B cells *in vivo*. In keeping with DNT expression in immature/transitional BM B cells, mtDNA was already reduced about ten-fold in resting splenic B cells of DNT mice (Figure 1e) and B cell activation by LPS was not able to increase the amount of mtDNA further, in contrast to WT B cells (Figure 1b, e).

**Figure 1.**
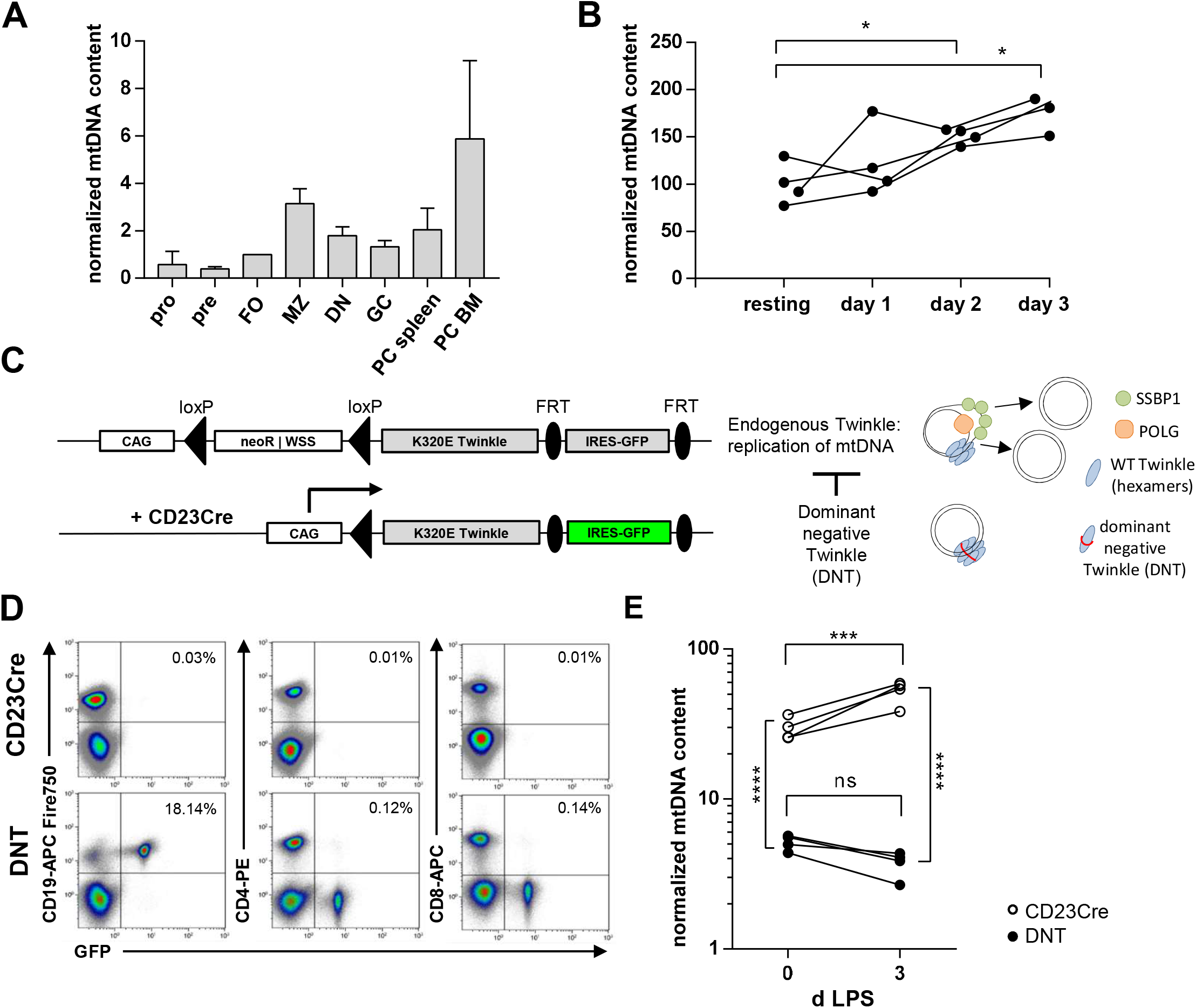
Expression of mtDNA and genetic inhibition of mtDNA replication in murine B cell subsets. A, Relative abundance of mtDNA encoded *16s rRNA* normalized to nuclear DNA with *hk2* as reference gene in sorted B cell subsets of SRBC immunized mice. n = 4, mean -/+ SEM. B, Splenic B cells were stimulated with LPS for 3 d. mtDNA abundance was assessed at given time points as described in A. C, Schematic of the construct encoding a dominant negative form of TWINKLE (K320E; DNT; (Baris et al., 2015) knocked into the *ROSA26* locus in the inactive conformation (upper) and after CRE mediated recombination (lower) with excised *neo*/Stop cassette (neo | WSS), activation of the composite CAG promoter (C, cytomegalovirus, A, chicken beta actin, G, rabbit beta globin) and expression of DNT and IRES-GFP. Expression of DNT impairs replication of mtDNA (right). D, Representative merged dot plots from splenic B cells from CD23CRE and DNT mice stained with antibodies against CD4, CD8 and CD19 and analyzed by flow cytometry. E, Splenic B cells from CD23CRE and DNT mice were stimulated with LPS for 3d and mtDNA was quantified on d0 and d3 as in A. Symbols represent individual mice. N=1; n=4; significance was calculated using 2-way ANOVA. Data for d3 are representative for 3 independent experiments.

### Replication of mtDNA in B cells is essential for humoral immunity

Since we observed GFP expression already in transitional B cells, we assessed B cell maturation in spleens of DNT mice with CD23CRE mice as controls (see representative FACS plots in Figure 2a-c). The total numbers of MZ B cells and plasmablasts/plasma cells were comparable and FO B cell numbers were even increased. The frequencies of GC B cells in Peyer’s patches were normal. Numbers of plasmablasts/plasma cells in the BM were also not altered (Figure 2d). However, the frequency of GFP^+^ MZ B cells was lower than in the FO B cell population (Figure 2e). More strikingly, the frequency of GFP expressing GC B cells was reduced to ∼20%, with a further decrease (down to ∼8%) in TACI^+^CD138^+^ BM plasmablasts/plasma cells. Plasmablasts/plasma cells in the spleen showed also a reduced GFP expression (∼25%). We conclude that DNT-expressing transitional B cells have a minor intrinsic disadvantage during MZ B cell development, which is consistent with the relatively high mtDNA amount in MZ B cells. In addition, DNT B cells have a strong disadvantage during the GC reaction that is even more pronounced in BM plasmablasts/plasma cells. The development of splenic plasmablasts/plasma cells, a heterogeneous population of short-lived, long lived, IgM expressing, and class-switched cells derived from the GC reaction and extrafollicular responses (Schuh et al., 2020) is also hampered by DNT expression. Unexpectedly, considering the normal numbers of plasma cells, serum abundance of class-switched Ig isotypes was reduced (Figure 2f) while IgM was not. A significant fraction of IgM is secreted by extrafollicular plasmablasts (Schuh et al., 2020). To determine how DNT-expression affects various plasma cell subtypes in the spleen and BM we analyzed plasma cell subsets in detail based on CD138, TACI, CD19 and B220 expression, namely the non-proliferating P3 cells (CD138^high^, TACI^high^, CD19^low^, B220^low^, BLIMP1^high^), the non-proliferating P2 cells (CD138^high^, TACI^high^, CD19^int^, B220^low^, BLIMP1^med^) and the proliferating P1 (CD138^high^, TACI^high^, CD19^int^, B220^int^, BLIMP1^low^) cells (Cossarizza et al., 2019; Pracht et al., 2017) (Figure 3a). Total cell numbers of P1, P2 and P3 plasma cells in spleen and BM were not affected by DNT expression (Figure 3b). Moreover, GFP expression was similar in B cells and P1 plasma cells in the spleen (76.5 % vs. 65.6 %) (Figure 3c). Interestingly, the P1 plasma cell population had a lower frequency of GFP expression in the BM than in the spleen (40.4% vs. 65.6%), suggesting that DNT expression disfavours the transition of plasma cells from the spleen to the BM. Strikingly, the frequency of GFP expressing cells dropped sharply in the non-proliferating P2 (spleen: 28.3%, BM: 8,9%) and even more so in P3 cells (spleen:10.1 %, BM: 3.1%) (Figure 3c). Hence, DNT-expression confers a disadvantage to developing plasma cells. Consequently, P2 and P3 plasma cells without the recombined DNT locus fill up the plasma cell compartments in spleen and BM. We conclude that a functional respiratory chain, which requires continuous replication of mtDNA, is necessary for the generation of plasma cells in the bone marrow. To challenge B cells, we immunized mice with sheep red blood cells (SRBC). Splenic GC formation as assessed by histology was severely impaired in SRBC immunized DNT mice (Figure 4a), pointing to a profound requirement of the mtRC for the GC reaction. Immunization of DNT mice with the TD antigen Nitrophenol-keyhole limpet hemocyanin (NP-KLH) revealed a ∼10-fold reduction of primary and a 4-5-fold reduction of secondary NP specific IgG responses (Figure 4b). Surprisingly, NP-KLH elicited NP-specific IgM was not reduced (Figure 4b) although DNT-expressing plasmablasts/plasma cells appear to have a specific developmental disadvantage as indicated by the homeostatic outcompetition of GFP^+^ cells (Figure 2e).

**Figure 2.**
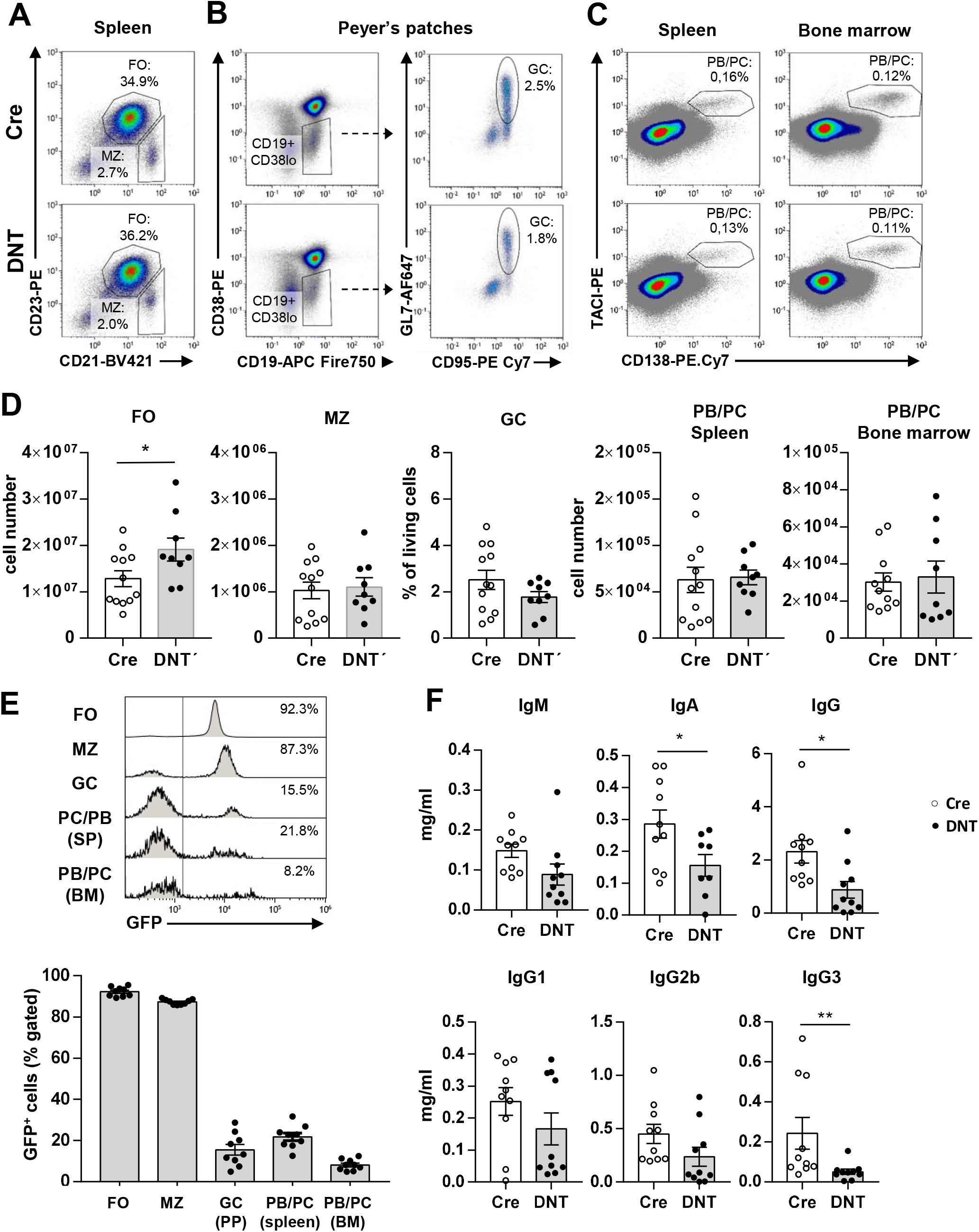
Quantification of B cell populations and serum antibodies in non-immunized DNT mice. A, Representative flow cytometry dot plots (merged from 3 mice) of splenic B cells pregated on singlets, viable lymphocytes and B cells (CD19^+^) from CD23CRE and DNT mice with anti CD21 and anti CD23 antibodies, B, Representative flow cytometric analysis of germinal center (GC) B cells in the peyer’s patches from CD23CRE and DNT mice pregated on singlets and viable lymphocytes stained with anti CD19, CD38, GL7 and CD95 antibodies. C, Representative flow cytometric analysis of plasma blasts and plasma cells defined as TACI^+^CD138^+^ from the spleen (left) and the bone marrow (right) from CD23CRE and DNT mice. D, Total cell numbers of follicular (FO) B cells, marginal zone (MZ) B cells, and plasma blasts/cells as well as frequencies of GC B cells of Peyer’s patches are shown in A-C. Data are presented as mean values. Symbols represent individual mice. N=3, n=2-4; significance was calculated using Mann-Whitney test. E, Representative flow cytometry dot plots (merged from 3 mice) of DNT/GFP expressing cells in indicated B cell populations of DNT mice and frequencies of GFP expressing cells in the indicated B cells populations of DNT mice. Data are shown as mean values. Symbols indicate individual mice. N=3, n=2-4; significance was calculated using 2-way ANOVA. F, Serum antibody concentrations of indicated isotypes from unimmunized CD23CRE and DNT mice (8-15 weeks old) were determined by ELISA. Data are shown as mean values. Symbols indicate individual mice. N=3, n=2-4; outliers excluded after Routs test; significance was calculated using Student’s t-test.

### Replication of mtDNA supports class switch recombination to IgG

It remained unclear which particular mechanisms were involved in the different NP-specific IgM and IgG responses. The decreased NP-specific IgG response could be due to the reduced GC reaction, reduced class switch recombination (CSR) or reduced GC derived plasma cell development, as already suggested in Figure 2e. Jang et al. proposed that activated B cells with a high mitochondrial content are more prone to undergo CSR (Jang et al., 2015). To determine whether mtRC activity is required for CSR we stimulated splenic B cells with LPS, anti CD40 antibody, IL-4, IL-5, transforming growth factor β (TGFβ) and retinoic acid (modified after (Chiu et al., 2019)). CSR to IgG was reduced by 40% while CSR to IgA was not affected as measured by the frequencies of IgA and IgG expressing B cells (Figure 4c,d). DNT expressing cells as indicated by GFP expression were only slightly underrepresented in the IgA^+^ fraction (Figure 4e, f). This result is consistent with the observation that homeostasis of existing IgA^+^ cells appears to be mediated through glycolysis (Kunisawa, 2017). We conclude that a functional mtRC is important for CSR to IgG.

### The development of antigen specific bone marrow plasma cells depends on OxPhos

Next, we addressed why NP-specific IgM responses were unaltered, but IgG responses were reduced in NP-KLH immunized DNT mice. Already in non-immunized mice, expression of DNT in GC B cells and in P2 and P3 plasma cells from spleen and BM, as determined by GFP expression, was significantly lower than in FO B cells (Figure 2e, 3b, c). We interpret these data as intercellular competition between B cells expressing a functional mtRC and those with GFP expression and reduced mtDNA. To show this directly we analyzed the development of antigen-specific plasma cells after the NP-KLH immunization using the CD138, TACI, CD19 and B220 staining applied in Figure 3 (Cossarizza et al., 2019; Pracht et al., 2017). Antigen specificity was determined by surface staining with NP-PE, which detects antigen-specific, surface IgM and IgA expressing plasma cells (Blanc et al., 2016). Immunization with NP-KLH clearly elicited NP-specific, CD138^high^ plasma cells in control and in DNT mice (Figure 5a). However, the cells binding surface NP-PE in DNT mice were hardly GFP positive (Figure 5a,b) as NP-PE staining and GFP expression were almost exclusive. In accordance with data from unimmunized animals (Figure 3), the frequencies of plasmablasts/plasma cells regardless of their proliferative state (Pracht et al., 2017) were unaltered but the frequencies of NP-binding plasma cells were reduced by ∼40% in immunized DNT mice (Figure 5c). Together, these data suggest that DNT-expressing plasmablasts are outcompeted during TD immune responses by NP-specific plasmablasts lacking transgenic DNT expression. To determine at which developmental stage DNT expressing plasmablasts are outcompeted, we analyzed GFP expression in P1, P2 and P3 plasma cells (Figure 5d, e). GFP expression was reduced in all the aforementioned subsets but the lowest GFP expression was detected in the P3 plasma compartment (2.2%). Together these data show that DNT expression interferes with the abundance of NP-specific plasma cells, whereas cells with intact mtDNA replication are not affected and show a relative expansion.

**Figure 3.**
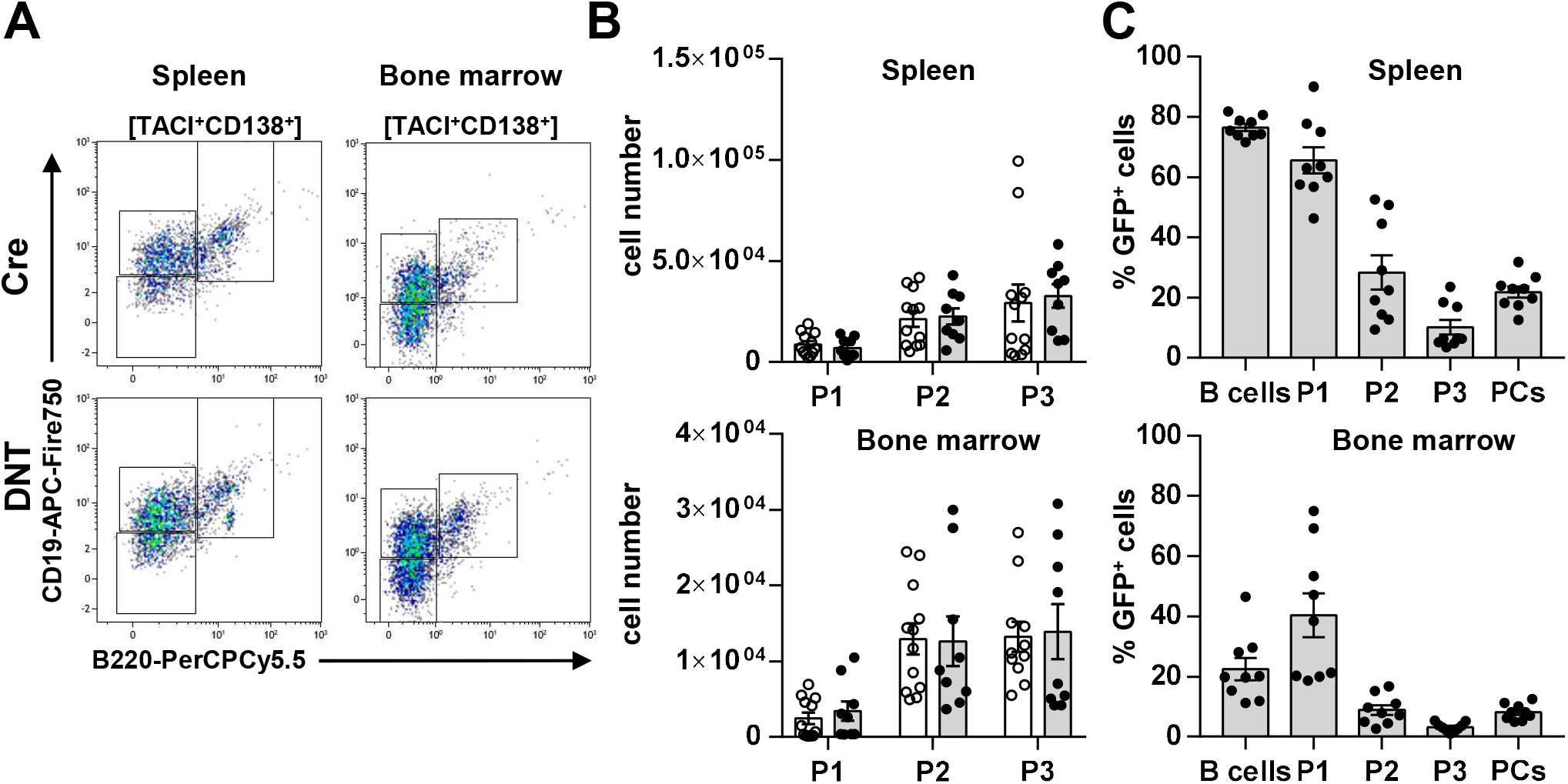
Characterization of plasma cell populations in non-immunized DNT mice. A, Representative flow cytometry dot plots of plasma cells (TACI^+^CD138^+^) (see Figure 2) from CD23CRE and DNT mice stained with anti CD19 and anti B220 antibodies, B, Total cell numbers of P1 cells (CD138^high^, TACI^high^, CD19^int^, B220^int^), P2 cells (CD138^high^, TACI^high^, CD19^int^, B220^low^) and P3 cells (CD138^high^, TACI^high^, CD19^low^, B220^low^) in the spleen and and bone marrow. Each dot represents one mouse, data are from 3 experiments. C, Frequencies of GFP expressing cells in B cells (CD19+, B220+), plasma cells (TACI^+^CD138^+^), P1 cells (CD138^high^, TACI^high^, CD19^int^, B220^int^), P2 cells (CD138^high^, TACI^high^, CD19^int^, B220^low^) and P3 cells (CD138^high^, TACI^high^, CD19^low^, B220^low^) in the spleen and bone marrow. Each dot represents one mouse, data are from 3 experiments.

### Replication of mtDNA sustains T cell-dependent and T cell-independent proliferation of B cells

The GC reaction, CSR and plasma cell differentiation all rely on sustained proliferation of activated B cells. Therefore, we determined whether and how prevention of mtDNA replication affects B cell proliferation, survival and plasmablast development *in vitro*. Survival of LPS activated B cells was not affected by DNT expression after 3 of activation (Figure 6a, b). To assess proliferation and plasmablast differentiation simultaneously we used a dye dilution assay and stained cells with the plasma cell marker CD138 (Nutt et al., 2011). This experiment showed that proliferation elicited by LPS as well as by CSR-inducing conditions (see Figure 4) was reduced by DNT expression (Figure 6c, d; Figure S3). To control this experiment intrinsically we took advantage of the fact that GFP expression of DNT B cells was not 100% (Figure S2b). This experiment revealed that DNT-expressing cells (green symbols, Figure S3b) accumulated less in divisions 5 and 6 under CSR inducing conditions and with LPS compared to B cells from the same culture that have not recombined (grey symbols, Figure S3b) or CD23CRE expressing B cells (black symbols, Figure S3b). However, we found DNT expressing LPS blasts that had divided 5 or 6 times (Figure 6 c, d, e; Figure S3b) but even those cells expressed less CD138 (Figure 6f). We conclude that DNT expression limits plasmablast development by limitation of proliferation and differentiation. To assess later time points we also investigated survival under CSR-inducing conditions because B cells live up to 9d under these conditions. However, in this setting, survival of DNT-expressing B cells dropped significantly on d7 and d9 and the surviving DNT-expressing cells displayed a markedly increased side scatter (Figure S4).

**Figure 4.**
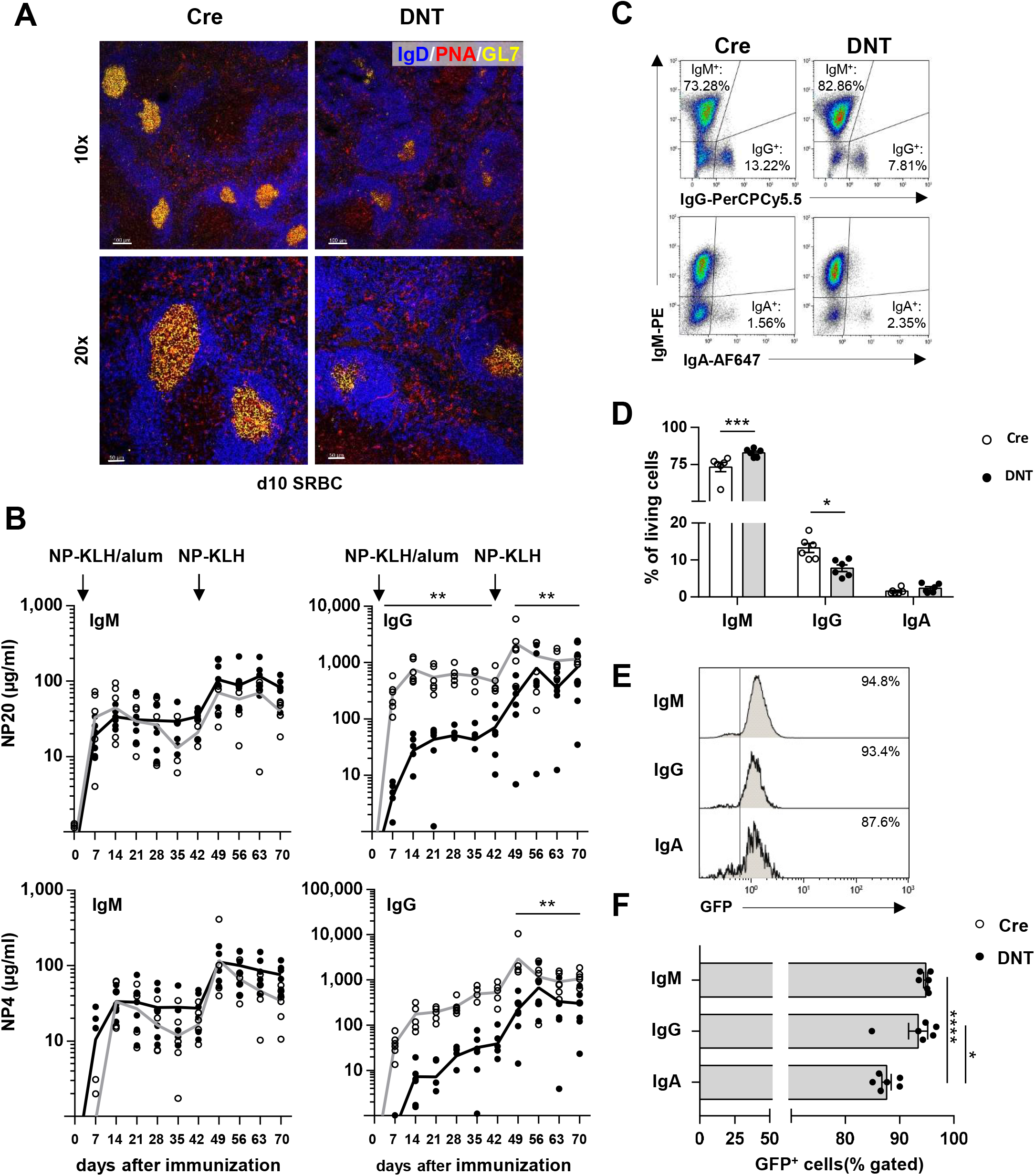
Impaired germinal center reactions, NP response and class switch recombination by DNT expression in B cells. A, CD23CRE and DNT mice were immunized with SRBC. Spleen sections were analyzed by histology and confocal microscopy. Upper panel: 10 x; lower panel 20 x; blue = IgD-AF647; green = GL7-PacificBlue; red = PNA-Rhodamine. Scale bars, 100 µm (top) and 50 μm (bottom). B, CD23CRE and DNT mice were immunized with NP-KLH in Alum *i.p.* and at d42 with NP-KLH in PBS *i.p.* Anti NP(20)- and NP(4)-specific serum antibody concentrations of indicated isotypes were determined by ELISA. The line represents the simple moving average; Symbols indicate individual mice. N=2, n=3; Significance was calculated using 2-way ANOVA of the area under curve. C, Splenic B cells from CD23CRE and DNT mice were stimulated with LPS, anti CD40 antibody, IL-4, IL-5, TGFβ and retinoic acid for 4d. Fixed B cells were analyzed by flow cytometry by intracellular staining of IgA, IgG and IgM. Representative dot blots are shown. D, Frequencies of isotype switched (IgA, IgG) or non-switched (IgM) B cells of stimulated (see C) B cells from CD23CRE and DNT mice. Data are presented as mean; Symbols indicate individual mice; N=2, n=3; Significance was calculated using 2-way ANOVA. E, Representative flow cytometric analysis of GFP expressing cells in indicated isotype switched B cell populations of DNT mice at d4 after stimulation with LPS, anti CD40 antibody, IL-4, IL-5, TGFβ and retinoic acid. F, Frequencies of GFP expressing cells in the indicated isotype switched B cells populations (see E) of DNT mice at d4 after stimulation with LPS, anti CD40 antibody, IL-4, IL-5, TGFβ and retinoic acid. Data are shown as mean values. Symbols indicate individual mice. N= 2, n=3; significance was calculated using one-way ANOVA.

**Figure 5.**
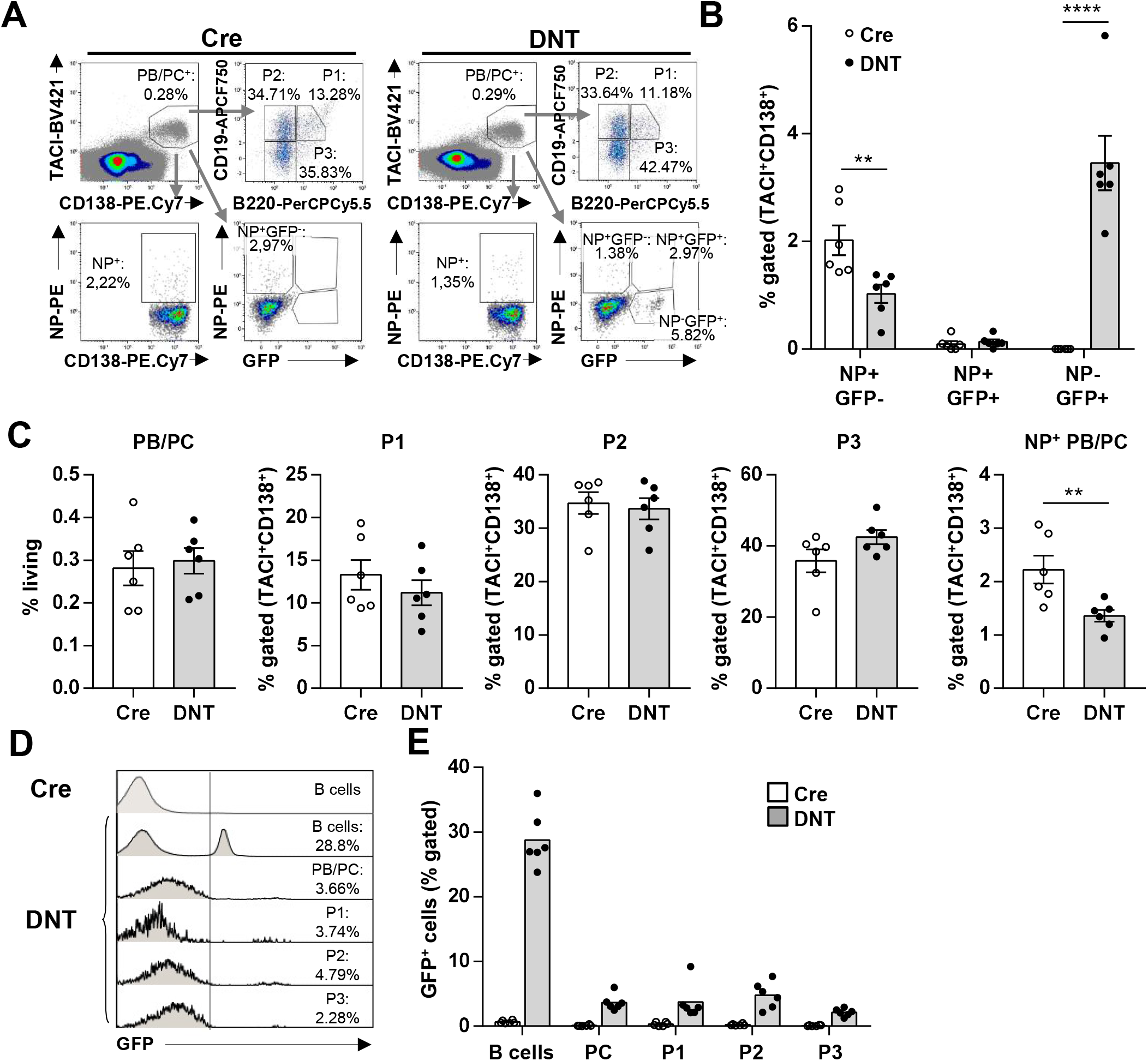
Decreased antigen specific plasma cells in NP-KLH immunized DNTWINKLE mice. A, CD23CRE and DNT mice were immunized with NP-KLH in Alum *i.p*. and challenged at d42 with NP-KLH *i.p.*. Plasma cells in the bone marrow were analyzed at d70 using anti CD138, TACI, CD19 and B220 antibodies to define P1, P2 and P3 populations (merged data of 3 mice from one experiment are shown). Antigen specific plasma cells were detected using surface staining with NP-PE. B, Frequencies of NP^-^GFP^+^, NP^+^GFP^+^ or NP^+^GFP^-^ cells in the TACI^+^ and CD138^+^ bone marrow cell population (shown in A). C, Frequencies of plasma blasts (PB) and plasma cells (PC), P1, P2, P3 and antigen specific plasma cells in the bone marrow as shown in A-B. Data shown as mean; Symbols indicate individual mice; N=2, n=3; Significance was calculated using 2-way ANOVA. D, Flow cytometric analysis of GFP expressing cells in the indicated B cells populations of DNT and CRE mice shown in A. E, Frequencies of GFP expressing cells in the indicated B cells populations of DNT and CRE mice shown in D. Data are shown as mean values. Symbols indicate individual mice. N= 2, n=3.

**Figure 6.**
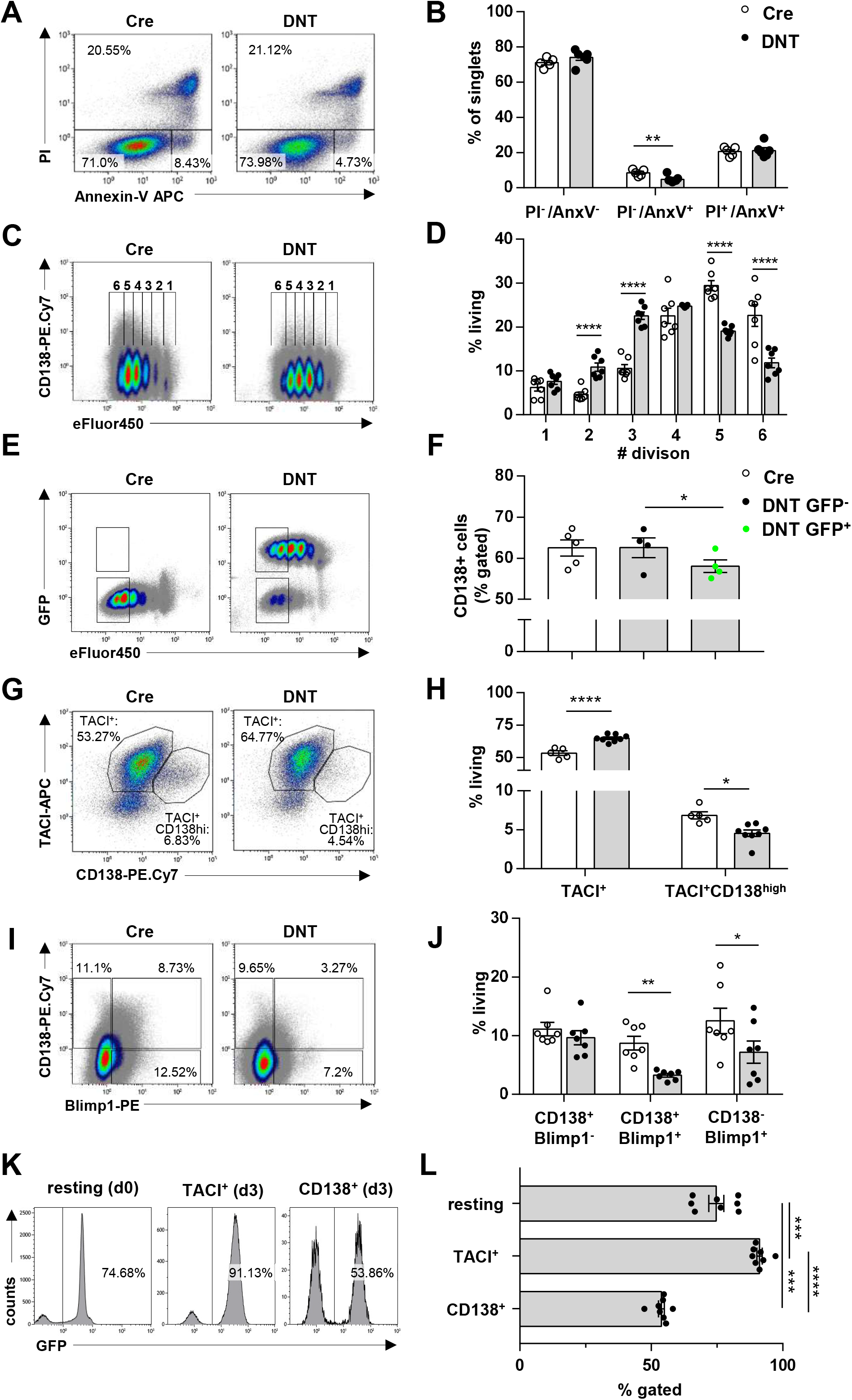
Comparable survival but decreased proliferation and plasma blast differentiation in LPS activated B cells from DNT mice. A, Splenic B cells from CD23CRE and DNT mice were stimulated with LPS for 3d, stained with propidium iodide (PI) and Annexin V and analyzed by flow cytometry. A representative, merged (3 mice per genotype) dot blot is shown. B, Frequencies of living (PI^-^, Annexin V^-^), apoptotic (PI^-^, Annexin V^+^) and necrotic (PI^-^, Annexin V^+^) cells on day 3 after LPS activation shown in A. Bars represent the mean, symbols represent individual mice. N=2; n=2-4; significance was calculated using 2-way ANOVA. C, Splenic B cells from CD23CRE and DNT mice were labelled with eFluor450, stimulated with LPS for 3d, stained with anti CD138 antibody and analyzed by flow cytometry. A representative, merged (3 mice per genotype) dot blot is shown. D, Frequencies of cell populations defined according to the number of cell divisions from CD23CRE and DNT mice on day 3 after LPS activation shown in C. Data are shown as mean values; Symbols represent individual mice. N=2; n=3; Significance was calculated using 2-way ANOVA. E, EFluor450 labelled splenic B cells from CD23CRE and DNT mice were stimulated with LPS for 3d, stained with anti CD138 antibodies and analyzed by flow cytometry. F, Frequencies of CD138 expressing cells in the GFP^+^ and GFP^-^ gates shown in E. Significance was assess by paired t-test. G, Splenic B cells from CD23CRE and DNT mice were stimulated with LPS for 3d, stained with anti CD138 and anti TACI antibodies and analyzed by flow cytometry. A representative, merged (3 mice per genotype) dot blot is shown. H, Frequencies of TACI^+^ and TACI^+^CD138^+^ cells shown in E. Data are shown as mean values; Symbols represent individual mice. N = 3; n= 2-4; Significance was calculated using 2-way ANOVA. I, Splenic B cells from CD23CRE and DNT mice were stimulated with LPS for 3d, fixed, permeabilized, stained with anti CD138 antibody on the surface and intracellular anti BLIMP1 antibodies and analyzed by flow cytometry. A representative, merged (3 mice per genotype) dot plot is shown. J, Frequencies of the CD138^+^, BLIMP1^+^ and BLIMP1^+^CD138^+^ cells shown in G. Data are shown as mean values; Symbols represent individual mice. N = 2; n= 2-4; Significance was calculated using 2-way ANOVA. K, Representative flow cytometric analysis of GFP positive, resting, TACI^+^, and TACI^+^CD138^+^ cells shown in E. L, Frequencies of GFP expressing cells in the indicated B cells populations of DNT mice shown in E. Data are shown as mean values. Symbols indicate individual mice. N = 2; n= 3; Significance was calculated using 2-way ANOVA.

### LPS-induced plasmablast development requires a functional mitochondrial respiratory chain

Although survival of LPS activated DNT B cells was not altered on d3, the frequency of CD138/TACI-expressing and CD138/BLIMP1 expressing plasmablasts was diminished (Figure 6g-j). In contrast, TACI expressing cells were proportionally increased (Figure 6g, h). In agreement with an accumulation of DNT-expressing plasmablasts in the TACI positive stage (Figure 6g, h), GFP-expressing cells were enriched at this stage but outcompeted by 40% at the TACI/CD138 positive stage (Figure 6k, l). In addition, immunoblots of day 3 LPS blasts showed that BLIMP1, XBP1 and IgM protein abundancies were lower while the amount of IRF4 was comparable (Figure 7a, b). Moreover, secreted IgM, IgG and IL-10 were also less abundant in DNT cell culture supernatants (Figure 7c, d). We conclude that DNT expression interferes with B cell intrinsic plasma cell differentiation but not with B cell activation elicited by LPS or LPS/anti-CD40/IL-4/IL-5/TGFβ/retinoic acid. We next examined cellular and mitochondrial morphology of resting and LPS-activated splenic B cells by electron microscopy (Figure 7e). Resting DNT-expressing splenic B cells having already 10-fold reduced mtDNA (see Figure 1e) did not contain less mitochondria than controls and their shape did also not differ grossly, but they were less electron-dense, indicating a looser cristae structure (Figure 7e, upper panel). This picture changed dramatically in LPS activated cells. We focused on activated B cells without obvious plasma cell morphology (compare Figure 7e, lower panel), keeping in mind that in DNT-cultures only roughly 60% of CD138-expressing plasmablasts were GFP positive (Figure 6j) which hindered the unambiguous identification of DNT-expressing plasmablasts. Activated B cells from CRE mice contained a dense ER and homogeneously electron-dense mitochondria while B cells from DNT mice were very heterogeneous for both parameters (not shown). However, the LPS-activated DNT B cells had few mitochondria that were extremely large and had a very low electron density. The cristae structure was almost completely lost (Figure 7e, lower panel). Thus, resting B cells, which are metabolically rather quiescent, can maintain almost normal mitochondrial morphology with 10-fold decreased mtDNA in contrast to 3d LPS activated B cells.

**Figure 7.**
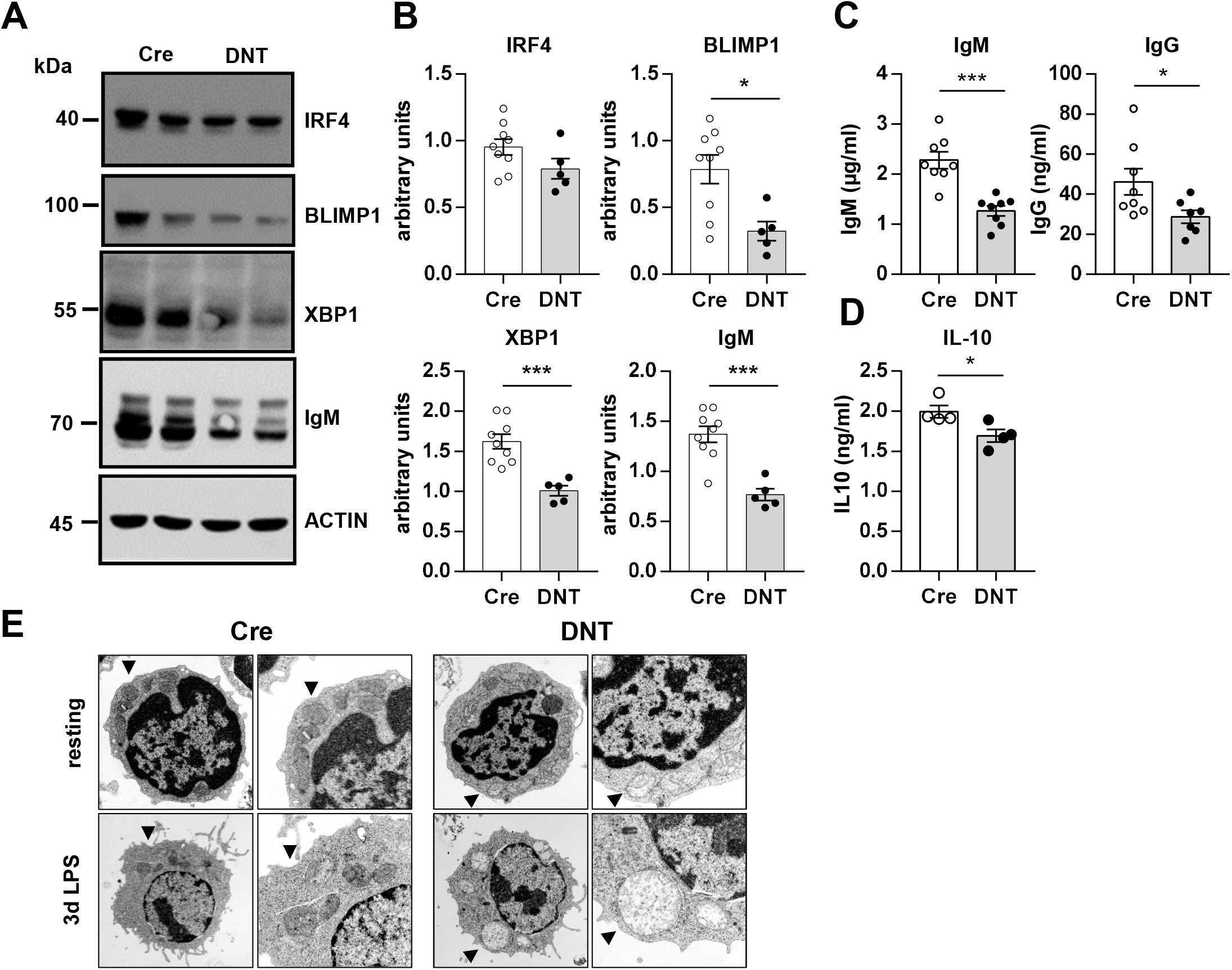
Decreased expression of plasma cell markers and antibody secretion in LPS activated B cells from DNT mice. A, Splenic B cells from CD23CRE and DNT mice were stimulated with LPS for 3d. Cell lysates were separated by 10% SDS-PAGE, transferred to nitrocellulose and stained with antibodies as indicated on the right. Molecular mass standards are shown on the left (kDa). A representative western blot showing lysates of each two mice is shown. B, Quantification of IRF4, BLIMP1, XBP1s and IgM protein expression relative to ACTIN of splenic B cells from CD23CRE and DNT mice stimulated with LPS for 3d. Symbols indicate individual mice. N=2, n = 2-4. Significance was calculated using 2-way ANOVA. C, Splenic B cells from CD23CRE and DNT mice were stimulated with LPS for 3d. Supernatants were analyzed by ELISA for the presence of IgM and IgG. Bars represent the mean ± SEM. Symbols indicate individual mice. N=2, n=4. Significance was calculated using Student’s t-test. D, Splenic B cells from CD23CRE and DNT mice were stimulated with LPS for 3d. Supernatants were analyzed by ELISA for the presence of IL10. Bars represent the mean ± SEM. Symbols indicate individual mice. N=1, n=4. Significance was calculated using t-test. E, Splenic B cells from CD23CRE and DNT mice were left unstimulated or stimulated with LPS for 3d, fixed, embedded and analyzed by transmission electron microscopy. Upper panel: resting cells (x 10.000, enlargements indicated by arrows: x 21.560), lower panel: blasts (x 6000, enlargements x 16.700)

### OxPhos is required for TCA flux in B cells

Next, we determined how reduced mtDNA affects oxidative phosphorylation and TCA metabolism during B cell activation and plasma cell differentiation (Figure 8). Extracellular flux analyses of d3 LPS cultures revealed that basal and maximal oxygen consumption as well as ATP production coupled to respiration were reduced in DNT B cells, while basal ECAR, glucose consumption and lactate secretion were enhanced (Figure 8a-d), revealing increased glycolysis in the presence of oxygen. The same experiment performed with cells activated for 6h showed that DNT-expressing cells performed OxPhos to a similar extent as controls although mtDNA was already depleted (see Figure 1) (Figure S5a). This suggested that mitochondrial function was still sufficient although loosening of cristae had already started (Figure 7e). In concordance with reduced OxPhos and ATP production linked to impaired respiration as measured by extracellular flux analyses (Figure 8b), we observed less ATP content in 3d LPS stimulated cells by luminometry (Figure 8, e). In contrast to the reduced mtDNA content already in resting splenic B cells but in accordance with the OxPhos measurements and mitochondrial morphology (Figure 1, Figure 7e and Figure S5a), ATP was not reduced in resting DNT B cells and even those cells were able to increase their intracellular ATP content upon activation (Figure 8e). This indicates compensatory pathways for ATP production, most probably glycolysis, which was increased in DNT-expressing B cells, but also hyper induction of mitochondrial biogenesis (Haumann et al., 2020).

**Figure 8.**
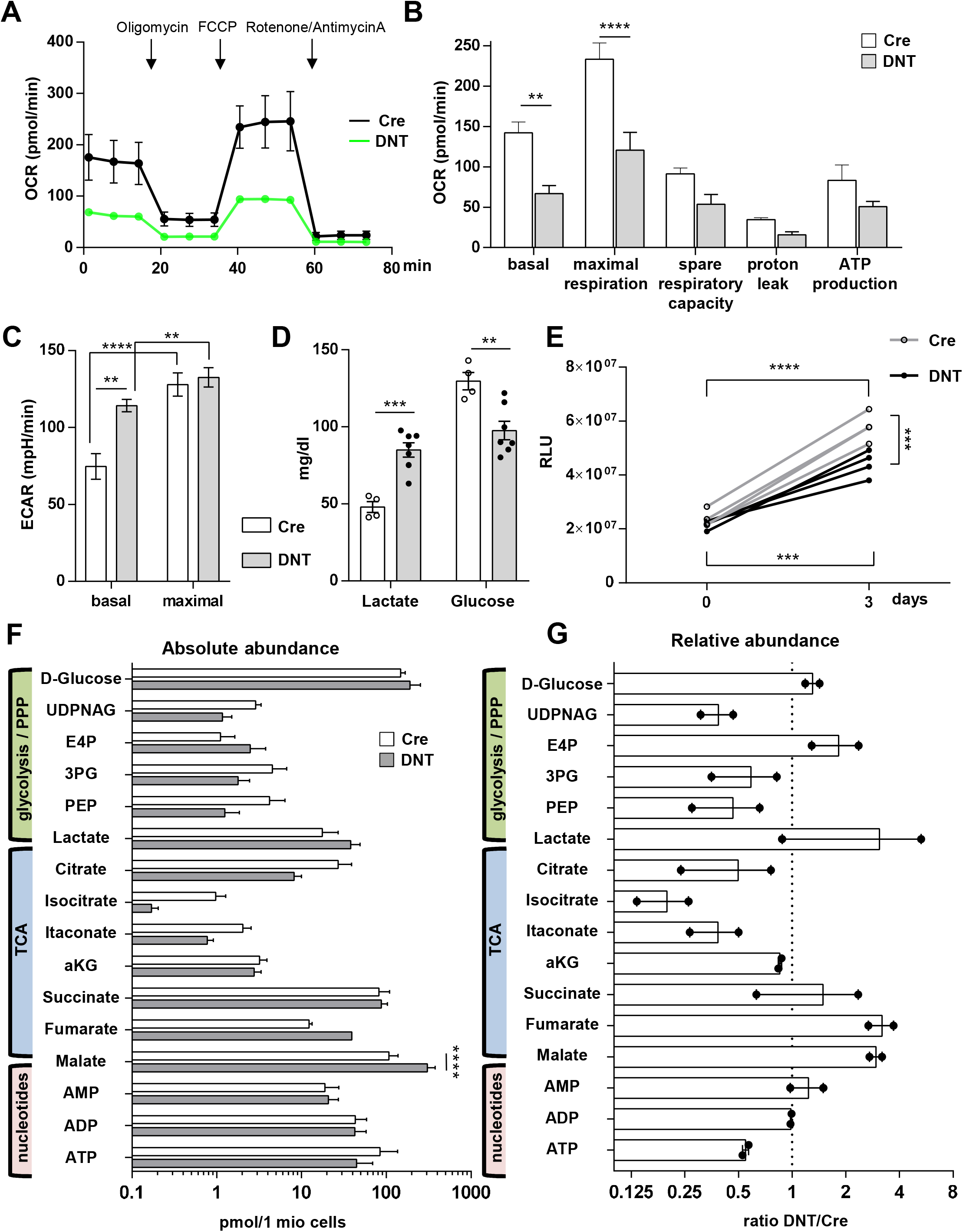
Impaired mtDNA replication in activated B cells shifts metabolism to glycolysis and changes intracellular metabolite concentrations. Splenic B cells from CD23CRE and DNT mice were analysed on day 3 after *in vitro* LPS activation. A, Representative extracellular flux analysis. The basal oxygen consumption rate (OCR) was measured before and after injection of oligomycin, FCCP and rotenone plus antimycin A. Symbols represent means of two mice, each of three replicate wells, and error bars are ± SEM. B, Calculated OCR values of basal and maximal respiration, spare respiratory capacity, and proton leak. Means ± SEM from N=2, n=2; values from different experiments were normalized using the mean of each genotype. Significance was calculated using 2-way ANOVA. C, Lactate and glucose concentrations in the supernatants on day 3 after LPS activation. Each dot represents one mouse, N=2, n=2-3. Significance was calculated using 2-way ANOVA. D, Luminometric detection of intracellular ATP levels of resting and day 3 activated B cells. RLU, relative light units. Symbols represent individual mice; Significance was calculated using 2-way ANOVA. E, HPLC/MS analysis of intracellular metabolites of FACS sorted B cells (DNT: viable GFP^+^, CD23CRE: viable). Data show the absolute abundance in pmol/10^6^ cells. Bars represent the mean with ± SEM; N=2, n=2; Significance was calculated using 2-way ANOVA. F, Relative abundance (DNT relative to CD23CRE) of intracellular metabolites shown in E. Bars at mean; Dotted line at 1; Symbols show the mean for each experiment; N=2, n=2.

The mtRC and the tricarboxylic acid cycle (TCA) are intertwined (Kennedy and Lehninger, 1949). Our data suggested that the mtRC is important for plasmablast differentiation during an anabolic phase in TACI^+^ cells. Hence, we measured glycolytic and TCA intermediates by liquid chromatography mass spectrometry (LC-MS) in d3 LPS cultures (Hofmann et al., 2011) (Figure 8 f, g; Figure S5b). The reduction of ATP (Figure 8b, e) was again confirmed by this approach. Moreover, in line with the extracellular glucose measurements (Figure 8d), intracellular glucose was slightly increased together with erythrose-4-phosphate (E4P). Phosphoenolpyruvate (PEP) and uridine diphosphate N-acetylglucosamine (UDPNAG) were less abundant. We found reduced metabolites upstream of succinate dehydrogenase (SDH) - citrate, isocitrate, itaconate - but an accumulation of the TCA metabolites fumarate and malate downstream of SDH, the only mtRC complex whose subunits are not encoded by mtDNA (Gustafsson et al., 2016) (Figure 8 f, g; Figure S5b, c). Altogether, these data suggest that B cells with a dysfunctional mtRC divert the TCA cycle and utilize glucose to supply the PPP similar to many tumor cells with mitochondrial dysfunction, thus explaining their initial ability to survive and sustain proliferation.

### The mitochondrial respiratory chain in plasmablasts coordinates lipid synthesis and MTOR activity

Succinate, fumarate and malate can inhibit prolyl hydroxylase domain (PHD)-containing enzymes that normally target the von Hippel-Lindau protein (vHL) which ubiquitinylates hypoxia inducible factor 1α (HIF1α), leading to its degradation (Harnoss et al., 2015; Martinez-Reyes and Chandel, 2020). Our metabolic analyses showed reduced ratios of α-ketoglutarate to fumarate and malate, indicating a pseudo-hypoxic state (Shanmugasundaram et al., 2014) (Figure S5b). In accordance, we observed a stabilization of HIF1α in DNT B cells at d3 of LPS stimulation under normoxic conditions (Figure 9a, b). The increased HIF1α protein abundance suggested increased reductive decarboxylation of α-ketoglutarate to citrate and the generation of long chain acylcarnitines (Xu et al., 2019), but our measurements indicated that citrate was not increased (Figure 8 f, g). Therefore, we reasoned that the driving force of the intact SDH complex II shifts the equilibrium of the TCA cycle away from citrate generated through reductive carboxylation of α-ketoglutarate towards the conversion of α-ketoglutarate to succinate (see Figure S5 for illustration). This would increase HIF1α abundance without concomitantly increasing citrate. Since HIF1α normally increases lipid synthesis in B cells via citrate (Xu et al., 2019), *de novo* lipid biosynthesis should be altered in DNT-expressing B cells. To investigate this issue, we performed lipidomic analyses of d3 LPS blasts (Kumar et al., 2015) (a representative mass spectrum is shown in Figure 9c). We observed a specific reduction of total phosphatidic acid (PA) (Figure 9d) but no major quantitative differences for phosphatidylglycerol (PG), phosphatidylinositol (PI), phosphatidylethanolamine (PE), phosphatidylcholine (PC) and phosphatidylserine (PS) (Figure 9d). Regarding the length and saturation of FA there were no obvious differences for PG. Nonetheless, there were remarkable differences for PA, PC, PI, PE and PS. Most dramatically, GPL species with short-chain and saturated or monounsaturated FAs such as 32:0, 34:0, 32:1 or 34:1 were reduced within the pools of PA, PI, PE and PS, with the exception of 30:0, 32:0 or 32:1 PC (Figure S6). Although we did not detect statistical differences for all longer FA chains, GPLs with long-chain, polyunsaturated ones such as 36:3-36:7 or 40:3-40:7 were increased continuously regardless of the headgroup (Figure S6). Since so many GPL species with polyunsaturated FAs - except within the class of PGs - were increased, we consider this effect cumulatively to be biologically relevant. While we detected higher levels of species with long-chain and polyunsaturated FAs within the pools of almost all GPL classes, PA was the only overall quantitatively reduced GPL class (Figure 9d). PA is a central precursor for many lipids, acts directly on MTOR (Menon et al., 2017) and is required for assembly and stability of MTOR complexes (Foster, 2013). MTOR in turn is required for plasma cell development (Jones et al., 2016). Hence, the reduced PA abundance in DNT-expressing B cells suggested that MTOR is activated less efficiently, which could explain the defective plasma cell development. In fact, phosphorylation of the ribosomal protein RPS6 (pRPS6), which is a substrate of the MTOR activated p70RPS6 kinase (Iwata et al., 2017), was reduced in DNT expressing LPS blasts at d2 and d3, but not at d1, while it was completely absent in resting cells (Figure 9a, e, f). Reduced MTOR activity could be a consequence of increased AMP activated kinase (AMPK) activity (Iwata et al., 2017;Tellier et al., 2016), which is activated by an increased AMP vs. ATP ratio. Corroborating our AMP and ATP measurements (Figure 8b, e-g), pAMPK was increased at d3 resulting in an elevated pAMPK vs. pRPS6 ratio in DNT-expressing B cells (Figure 9b, Figure S7). To assess MTOR activity kinetically we tracked pRPS6 over 3d of LPS activation (Figure 9e). Consistent with the immunoblot analyses (Figure 9a), flow cytometry revealed that RPS6 was less phosphorylated in 48h and 72h LPS activated DNT B cells (Figure 9e, f). Together, our data suggest that elevated AMPK activity and/or reduced PA abundance suppress MTOR in activated B cells.

**Figure 9.**
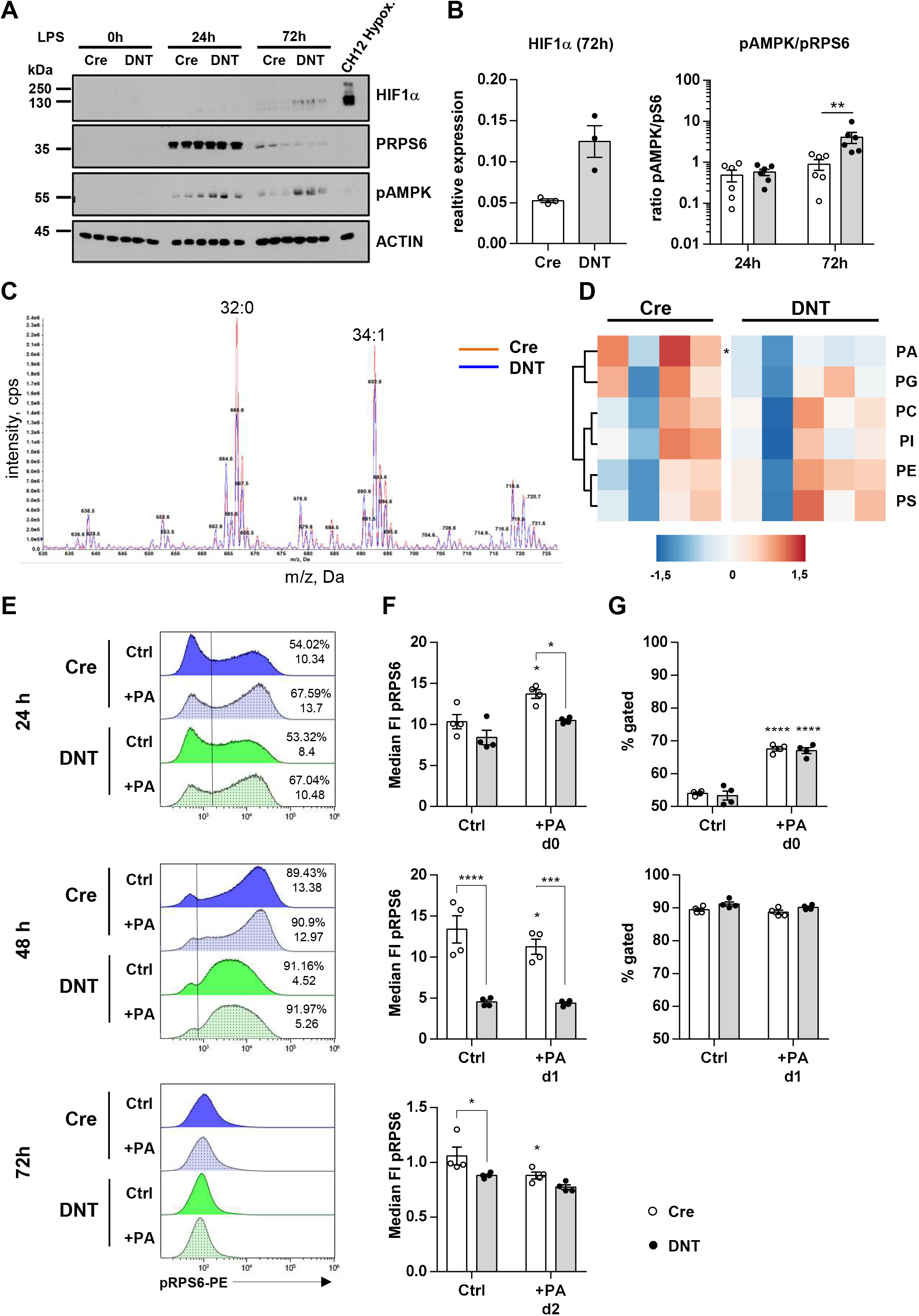
Oxidative phosphorylation controls MTOR activity in early plasmablasts via phosphatidic acid. A, Splenic B cells from CD23CRE and DNT mice were stimulated with LPS for 0, 24 h or 72 h. Cell lysates were separated by 10% SDS-PAGE, transferred to nitrocellulose and stained with antibodies as indicated on the right. Molecular mass standards are shown on the left (kDa). B, Quantification of HIF1α and pAMPK expression relative to Actin and of pAMPK relative to pRPS6. C, Splenic B cells from CD23CRE and DNT mice were stimulated with LPS for 3d. Glycerophospholipids of FACS sorted B cells (DNT: viable GFP^+^, CD23CRE: viable) were analyzed via direct infusion MS/MS (*Shotgun Lipidomics*). A representative neutral loss mass spectrum of phosphatidic acid species from Cre and DNT B cells (overlay) is shown, with small numbers indicating masses of the [M+NH4]^+^ precursor ions and large numbers indicating the total number of carbon atoms and the total number of double bonds within the two fatty acyl chains (two representative species, 32:0 and 34:1, are shown). D, The absolute abundance (nMol/mg protein) of phosphatidic acid (PA), phosphatidylglycerol (PG), phosphatidylinositol (PI), phosphatidylcholine (PC), phosphatidylethanolamine (PE) and phosphatidylserine (PS) is depicted as heatmap. Each symbol represents one mouse. One experiment out of two with identical results is shown. Significance was calculated using 2-way ANOVA. E, Splenic B cells from CD23CRE and DNT mice were stimulated with LPS in the presence or absence of phosphatidic acid (PA) liposomes and analyzed after 24 h (upper), 48 h (middle) and 72 h (please note different y-axis scaling) by flow cytometry. Representative histograms of pRPS6 expression are shown. Numbers indicate frequencies of pRPS6-expressing cells and the median fluorescence intensity of the positive cells F, The median pRPS6 fluorescence of is presented as median -/+ SEM. Symbols indicate individual mice; N=1, n=4. G, Frequencies of pRPS6 positive cells shown in E. Significance was calculated using 2-way ANOVA. The significance symbols (*) placed on top of the columns show the difference compared to the untreated controls of the respective genotype while the brackets indicate differences between the genotypes.

### Phosphatidic acid increases OxPhos-dependent MTOR activity in LPS activated B cells

Consequently, we tested the hypothesis that the lower abundance of PA in d3 DNT LPS blasts is responsible for suppressed MTOR activity. To this end, we added PA liposomes to LPS activated DNT blasts (Figure 9e-g; Figure S8). Addition of palmitic-acid-containing PA (32:0), which was reduced the most (Figure 9c, d; Figure S6), for 24h to LPS activated B cells increased the frequency of pRPS6 expressing cells in DNT B cells and controls similarly after 24h (Figure 9e-g). Nevertheless, there was no significant PA induced phosphorylation of RPS6 in DNT expressing B cells, showing that PA mediated mTOR activation requires a functional mtRC. PRPS6 increased further after 48h in LPS activated control B cells independently from PA addition but declined in DNT expressing B cells. After 72h PRPS6 also declined in the control cells (Figure 9a, e-g). These data show 1) that MTOR activity is only transiently elevated in LPS activated B cells, 2) that induction of MTOR via PA is only effective within the first 24 of B cell activation and 3) that induction of MTOR via PA in B cells depends on OxPhos. We propose an integrated model featuring a positive feedback loop between mitochondria and the nucleus that controls plasma cell generation (Figure 10). OxPhos, which requires continuous replication and transcription of mtDNA, drives the TCA cycle to generate PA, leading to MTOR activation and BLIMP1 induction. BLIMP1 then induces expression of OxPhos genes encoded in the nucleus, further increasing OxPhos in a progressive manner.

**Figure 10.**
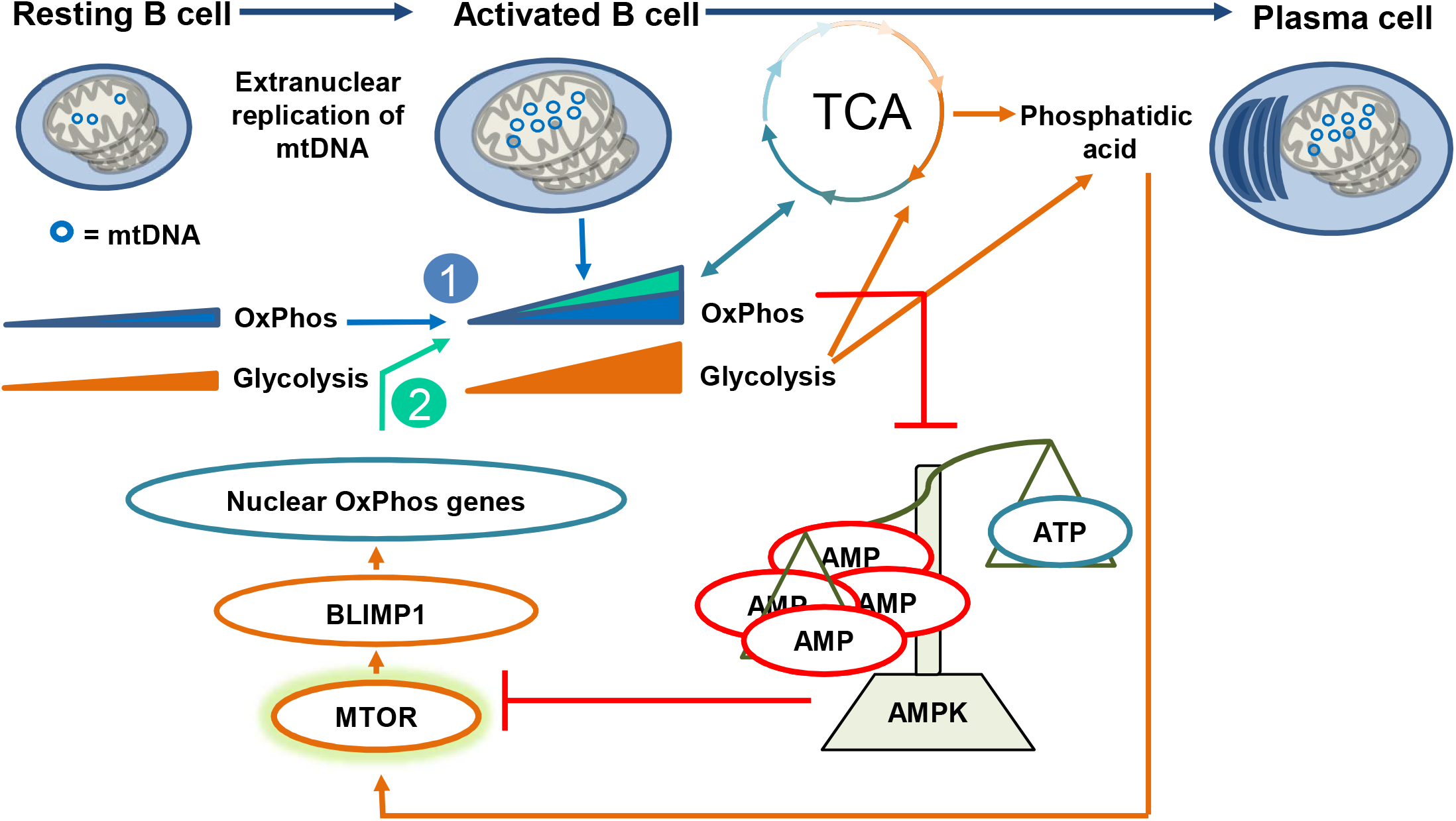
Integrated model of OxPhos controlled plasma cell development. Integrated model of OxPhos controlled plasma cell development showing that LPS-activated B cells replicate mitochondrial DNA which is essential for OxPhos and flux of the TCA cycle. A positive feedback loop between OxPhos and BLIMP1 expression controls plasma cell generation: the initial mtDNA dependent increase in OxPhos (1) drives the TCA cycle to generate phosphatidic acid, to activate MTOR and to inhibit AMPK which itself blocks MTOR in a AMP/ATP ratio-dependent manner. When OxPhos is active, the TCA cycle fluxes, MTOR is active and BLIMP1 is expressed (Gaudette et al., 2020; Jones et al., 2016). BLIMP1 then fosters up-regulation of nuclear OxPhos genes (2) and MTOR activity (Price et al., 2018; Tellier et al., 2016). Combined upregulation of mtDNA encoded essential mtRC subunits and nuclear mtRC components ensures full OxPhos activity which prevents AMPK activation and preserves MTOR function.

## Discussion

Here we describe selection checkpoints for mitochondrial function during B cell maturation, activation and plasma cell differentiation using depletion of mtDNA via CD23CRE-mediated expression of a dominant negative version of the mitochondrial helicase TWINKLE (DNT) coupled to IRES-GFP (Baris et al., 2015; Holzer et al., 2019; Weiland et al., 2018). The original human TWINKLE mutant, K319E, causes severe neurological problems (Hudson et al., 2005) but no data regarding humoral immunity are available. We show that MZ B cells and plasma cells contain the highest copy number of mtDNA within the B cell lineage, and that B cell activation is accompanied by increasing mtDNA. In line, the GC reaction, CSR, plasmablast and plasma cell development depend on sufficient copies of mtDNA. MZ B cell development was only slightly affected. Proportional to the highest mtDNA copy number plasma cells were most vulnerable to mtDNA depletion. In the plasma cell population as well as in the GC B cell population, GFP expressing cells become outcompeted by B cells that had escaped CD23CRE mediated excision of the STOP cassette. With regard to FO B cells our data support the finding that B cells do not require optimal expression of mitochondrial complex I, III, and IV proteins for their development (Milasta et al., 2016). In contrast to the work of Milasta et al., who conditionally deleted AIF1 (Milasta et al., 2016), we show that plasma cell development and function do depend on OxPhos. This contrast may be explained similar to our findings by the speculation that some B cells may also have escaped genetic recombination of the AIF1 locus (Milasta et al., 2016) and filled up the plasma cell compartment. We propose that the relative amount of mtDNA predicts the dependence of a given subpopulation on mtDNA, i.e. OxPhos activity. Hence, quantifying the mtDNA content of human B cell populations in health and disease will be important. Our data indicate that DNT expression by CD23CRE occurs in transitional B cells and consequently we found reduced mtDNA in resting FO B cells but no effect on their development. The amount of mtDNA does not increase in GC B cells albeit those cells as well as resting FO B cells are supported by mitochondrial FA metabolism (Caro-Maldonado et al., 2014; Weisel et al., 2020). Since GC B cells are the most rapidly dividing cells in the mammalian body (MacLennan, 1994) we conclude that cell proliferation in GC is accompanied by replication of mtDNA to ensure that the mtDNA content remains at least constant. While there are conflicting data concerning the role of hypoxia and OxPhos within the GC (Boothby and Rickert, 2017; Jellusova et al., 2017; Weisel et al., 2020), it should be noted that GC B cells are a heterogeneous population (Ise et al., 2018; Victora et al., 2010). Different metabolic requirements feeding back on one another may be required for GC B cell entry, progression, and exit (Ersching et al., 2017). We have not yet determined at which GC stage mtDNA replication is most important. Nonetheless, our results show that mitochondrial fitness determined by a functional respiratory chain, which requires continuous replication and translation of mtDNA, constitutes a bottleneck of B cell selection within the GC, by controlling GC B cell expansion and generation of antigen-specific IgG secreting plasma cells.

Unexpectedly, we found that DNT-expressing B cells proliferate initially well and show even slightly increased survival rates when activated by LPS or LPS/CD40/IL-4/IL-5/TGFβ/retinoic for 3 and 4 days although mtDNA copy number was ∼10 fold reduced and the mitochondrial cristae structure appeared already less dense. This was surprisingly paralleled by normal OxPhos and unaltered MTOR activity at d1 but less activity at d2 and d3. These data are consistent with reciprocally increased AMPK activity since AMPK inhibits MTOR and promotes mitochondrial function and quality in B cells (Brookens et al., 2020). Moreover, the reduced AMP/ATP ratio observed in the metabolomic analyses is consistent with increased AMPK activity. We conclude that one important function of OxPhos up-regulation in proliferating B cells is suppression of AMPK activity. These data are in accordance with original proposals linking the TCA, AMPK and MTOR (Tokunaga et al., 2004) and can indirectly also explain the effects on CSR and the GC reaction, both of which are controlled by MTOR and sustained proliferation (Chiu et al., 2019; Ersching et al., 2017; Keating et al., 2013; MacLennan, 1994; Stavnezer et al., 2008; Zhang et al., 2013). In line, AMPK has previously been shown to control the GC reaction (Lee et al., 2017; Waters et al., 2019). Our data are also consistent with DNT-mediated activation of AMPK in primary cultured chondrocytes (Holzer et al., 2019). Of note, AMPK activity is associated with aged, self-reactive and pro-inflammatory, CD21^-^CD23^-^ double negative (DN) B cells (Frasca et al., 2019). Along with the reduced IL-10 secretion we observed after LPS stimulation this might indicate a pro-inflammatory effect of DNT expression in B cells, an issue that deserves further investigation. Jang et al. have proposed that increased mitochondrial mass and ROS in activated B cells support CSR (Jang et al., 2015). Counterintuitively, Haumann et al. showed recently that mtDNA mutations induce a retrograde signaling pathway that actually increases mitochondrial biogenesis, thereby, increasing metabolic fitness and the tumourigenic potential of Reed-Sternberg cells (Haumann et al., 2020). One of the retrograde pathways inducing mitochondrial biogenesis (Haumann et al., 2020) could be the catabolic AMPK pathway (Brookens et al., 2020) that is elevated in DNT expressing B cells.

An exceptional advantage of our experimental model is the circumvention of unspecific and toxic inhibitors of OxPhos or glycolysis *in vivo*. This enabled us to analyze TD immunizations, revealing a very strong B cell intrinsic requirement of OxPhos during the TD NP-specific IgG response. The reduced basal IgG and the reduced immunization induced NP-specific IgG may represent a cumulative consequence of crippled GC reactions and CSR. Since we observed a drop in GFP – i.e. DNT – expression between the GC stage and BM plasma cells we propose a metabolic checkpoint during post-GC plasma cell development and/or survival. This proposal is supported by our *in vitro* data, albeit those were obtained in short-term cultures with TI stimulation. Additional strong support for the concept of a disadvantage of B cells with reduced OxPhos during TD-induced plasma cell generation is provided by the fact that GFP-expressing cells are excluded from the population of NP-specific plasma cells. We explain this by the expansion of GFP negative, NP-specific plasma cells identified by TACI/CD138 staining (Pracht et al., 2017) in NP-KLH immunized DNT mice. Like in classical GC defects such as during CD40/CD40L or activation-induced deaminase (AID) deficiency, the IgM expressing plasma cells, in our setting WT cells, may over-compensate (Durandy and Honjo, 2001) and may even secrete more Ab. Along this line, oligomycin, an inhibitor of ATP synthase, has been shown to reduce Ab secretion in LPS blasts (Price et al., 2018), albeit the concentrations that had been used are unclear. Another factor that may come into play is Ig catabolism, which is proportional to Ig abundance (Morell et al., 1970; Waldmann and Strober, 1969). Ab half-life may possibly be prolonged in DNTxCD23CRE mice.

Price et al. have revealed a progressive up-regulation of OxPhos accompanying plasma cell generation towards the TI antigen LPS through BLIMP1-controlled expression of OxPhos genes (Price et al., 2018). In contrast, other groups have shown that BLIMP1 reduces mitochondrial mass and mitochondrial reactive oxygen species (Jang et al., 2015). The survival of plasma cells in the BM, defined as CD138^+^B220^low^ (Lam et al., 2016), appears to require mitochondrial pyruvate import to prevent bioenergetic crises. Our data support the model that the generation BM plasma cells (compare GFP expression between plasma cell subsets from spleen and BM), require a functional mtRC fueled by the TCA cycle. Pyruvate is essential to generate citrate and consequently lipids. Based on the relative distribution of TCA metabolites in DNT expressing LPS blasts we hypothesized that the reduced citrate affects LPAAT pathway (Foster et al., 2014). In that sense, the reduced OxPhos in LPS activated B cells lowered the total abundance of PA, in particular PA species with saturated FAs, and shifted the overall equilibrium to GPLs with polyunsaturated fatty acyl chains. This can be expected to alter membrane dynamics and may explain the large side scatter of late DNT blasts (Tokumasu et al., 2009). One reason for decreased saturated GPLs could be facilitated β-oxidation (Schulz, 2008), a process that is also supported by AMPK (Garcia and Shaw, 2017). While addition of PA was able to enhance MTOR activity within the first 24h after activation, it did not at later timepoints. The effect of PA may depend on the cellular differentiation status, for instance, at later timepoints activated AMPK could restrict the PA effect. We used saturated palmitic-acid-containing PA (32:0) because this species was reduced the most in LPS-elicited DNT-blasts. Others have reported that only PA containing one unsaturated FA can activate MTOR (Yoon et al., 2015). The saturated PA may have been effective in our hands because we have used primary mouse B cells instead of transformed and adherent human cells. It has been proposed that the responsiveness of MTOR to PA evolved to sense lipid precursors for membrane biosynthesis prior to doubling the mass of a cell and cell division (Foster, 2013). We envision a similar function for MTOR in B cells. This would be in analogy to its function in T cell priming (Angela et al., 2016) and with the observation that MTOR is required during early steps of plasma cell differentiation by priming the endoplasmic reticulum for increased protein synthesis (Gaudette et al., 2020). In accordance, we observed numerous alterations in the lipid composition of DNT-expressing B cells, explaining why PA activates mTOR better in B cells with intact OxPhos. Although we are far from fully understanding the biological significance of the complex alterations in phospholipids, two issues deserve further investigation: Firstly, and as already discussed, the precise lipid-mediated mechanism of the regulation of MTOR activity in B cells. Secondly, the regulation of mitochondrial fission by Dynamin-like protein 1 (DRP1), which binds simultaneously to the polar headgroup of PA and to the acyl chains of other membrane phospholipids (Adachi et al., 2016). Indeed, mitochondria were enlarged in the DNT-expressing LPS blasts and it is tempting to speculate that this is due to missing action of DRP1. Our data obtained in LPS blasts, which are short-lived, may not completely reflect the metabolism of long-lived plasma cells but we clearly observe similar effects *in vitro* and during TD immune responses *in vivo*. Unless it is feasible to obtain sufficient numbers of long-lived plasma cells, for instance through *in vitro* cultivation systems (Cocco et al., 2012; Cornelis et al., 2020), LPS blasts serve to establish working hypotheses.

Many tumour cells feature defective mitochondrial respiration and rely on reductive carboxylation of glutamate to produce citrate (Mullen et al., 2011). Unlike tumour cells where the mtRC complexes are stochastically mutated, the B cells expressing DNT do have a fully functional complex II, the SDH complex, because no subunit of SDH is encoded by mtDNA (Gustafsson et al., 2016). We envision that the intact complex II is the driving enzymatic reaction in the TCA cycle of DNT B cells due to loss-of-function mutations in the other complexes. The shifts the equilibrium to more downstream (fumarate, malate) and less upstream metabolites (citrate, isocitrate, itaconate, α-ketoglutarate). Henceforth, less citrate can be used for FA synthesis and PA generation, the ratio of α-ketoglutarate to fumarate and malate is lowered in DNT B cells, which disfavors PHD activation (for review see (Martinez-Reyes and Chandel, 2020) and causes stabilization of HIF1α in DNT LPS blasts at d3. Although we have not shown this directly, our results are consistent with a role of glutamine in supporting mitochondrial respiration and anaplerotic reactions in the TCA cycle of plasmablasts (Lam et al., 2018).

Essentially, we provide a causal relation between OxPhos and plasma cell differentiation *in vivo* and in primary B cell cultures *in vitro* and place OxPhos mediated generation of TCA intermediates upstream of BLIMP1 and XBP1. This proposal stands in contrast to data showing that hypoxia increases IgG1 class switch and plasma cell differentiation (Abbott et al., 2016). The discrepancy may be due to the fact that our concept is based on data obtained in a B cell intrinsic manner. We propose a positive feedback loop between mtDNA controlled OxPhos and BLIMP1 mediated expression of nuclear OxPhos genes that ensures selection for mitochondrial fitness during plasma cell generation (Figure 10): the initial increase in OxPhos (Caro-Maldonado et al., 2014) drives the TCA cycle to generate PA and ATP, leading to inhibition of AMPK, activation of MTOR and subsequent BLIMP1 expression (Brookens et al., 2020; Gaudette et al., 2020; Jones et al., 2016). This occurs in activated, TACI^+^ B cells not yet expressing CD138. The higher the TCA cycle flux is, the more PA is generated, and the more BLIMP1 is expressed. BLIMP1 induces expression of OxPhos genes encoded in the nucleus, further increasing OxPhos in a progressive manner (Price et al., 2018). This model could also explain how BLIMP1 is able to sustain MTOR activity (Tellier et al., 2016), namely via OxPhos and the TCA cycle.

In conclusion, we identify here checkpoints for mitochondrial fitness during B cell activation and plasma cell differentiation. Functional OxPhos, which requires the continuous replication and transcription of mtDNA, prevents AMPK activation, and supplies the cell with TCA intermediates needed for MTOR activity and humoral immunity.

## Abbreviations

Ab: antibody
AMPK: Adenosine monophosphate activated kinase
BCR: B cell receptor
BM: bone marrow
CSR: class switch recombination
ETC: electron transport chain
FA: fatty acid
FAD: flavine adenine dinucleotide
GC: germinal center
G3P: Glyceraldehyde-3-phosphate
GPL: glycerophospholipid
IMM: inner mitochondrial membrane
IRF4: Interferon regulatory factor 4
LC-MS: liquid chromatography mass spectrometry
LPA: lysophosphatidic acid
LPAAT: Lysophosphatidic acid acyltransferase
MZ: marginal zone
mtDNA: mitochondrial DNA
mtRC: mitochondrial respiratory chain
NAD: nicotinamide adenine dinucleotide
NP-KLH: Nitrophenol keyhole limpet hemocyanin
PEP: phosphoenolpyruvate
PA: phosphatidic acid
PC: phosphatidylcholine
PE: phosphatidylethanolamine
PG: phosphatidylglycerol
PI: phosphoinositol
PS: phoshphatidylserine
PI3K: phosphoinositol-3-kinase
pRPS6: phosphorylated ribosomal protein RPS6
SRBC: sheep red blood cell
SHM: somatic hypermutation
SDH: Succinatedehydrogenase
TCA: tricarboxylic acid cycle
TD: T-cell dependent
TI: T-cell independent
UDPNAG: Uridine Diphosphate N-acetylglucosamine
vHL: von Hippel-Lindau protein

## Acknowledgements

This work was funded by the Deutsche Forschungsgemeinschaft (DFG; transregional collaborative research grant TRR130 and DFG Research training grant 1660, to D.M.). O.R.B. and R.J.W. were supported by the Deutsche Forschungsgemeinschaft (DFG; SFB 829/A14 and Cologne Excellence Cluster on Cellular Stress Responses in Aging-associated Diseases – CECAD).

## Declaration of interests

The authors declare no competing financial interests.

## Author contributions

S-U., O.R.B., F.G., U.S-S., J.H., S.B., W.S., T.S. performed experiments. S.U. and D.M. designed the study. S.U., F.G., U.S-S., J.H., S.B., R.W. and D.M. analyzed data. S.R.S. performed bioinformatics analyses. S.U., R.W. and D.M. wrote the paper. D.Mo., U.S., K.C. and R.W. provided intellectual and infrastructural help.

## KEY RESOURCES TABLE

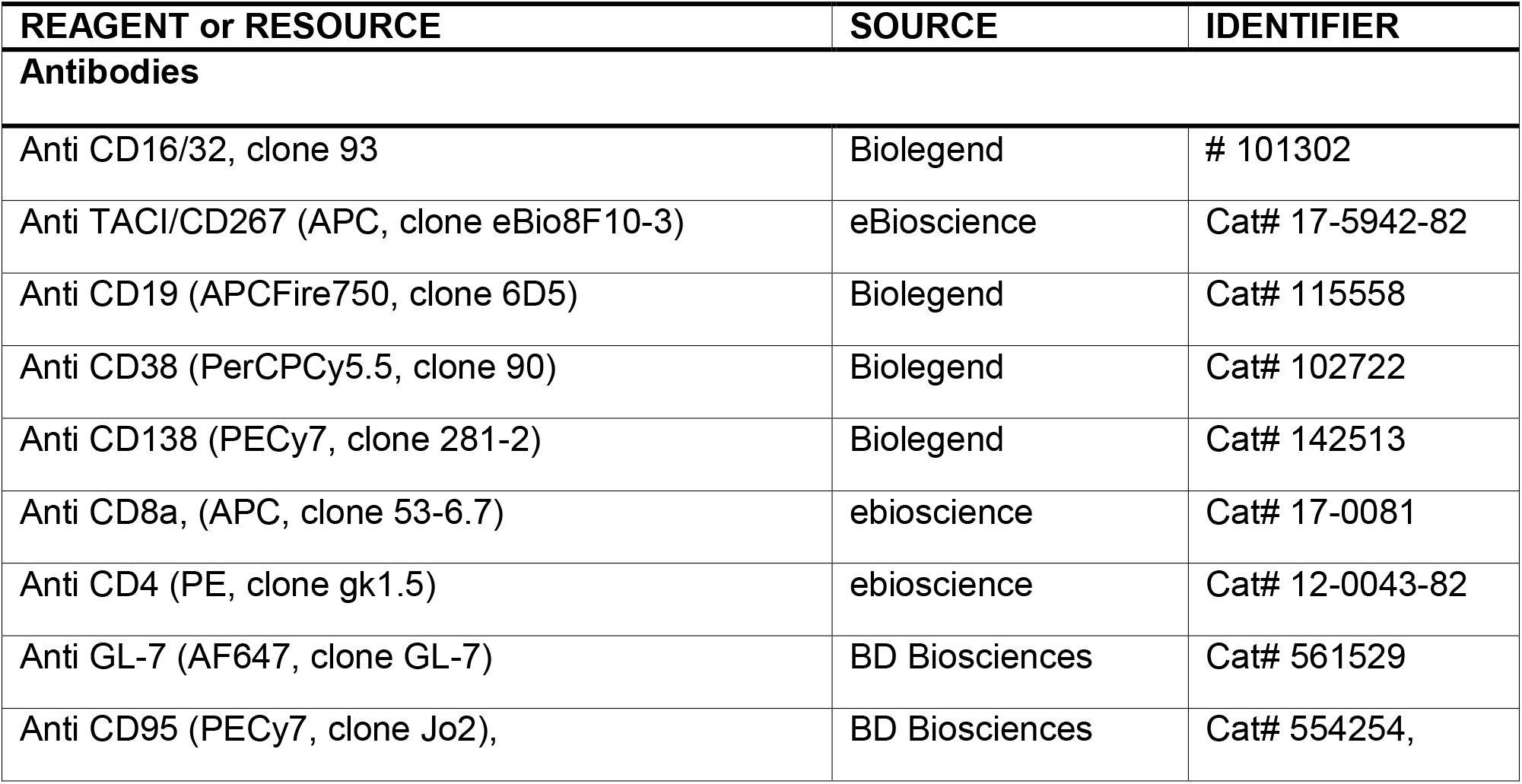
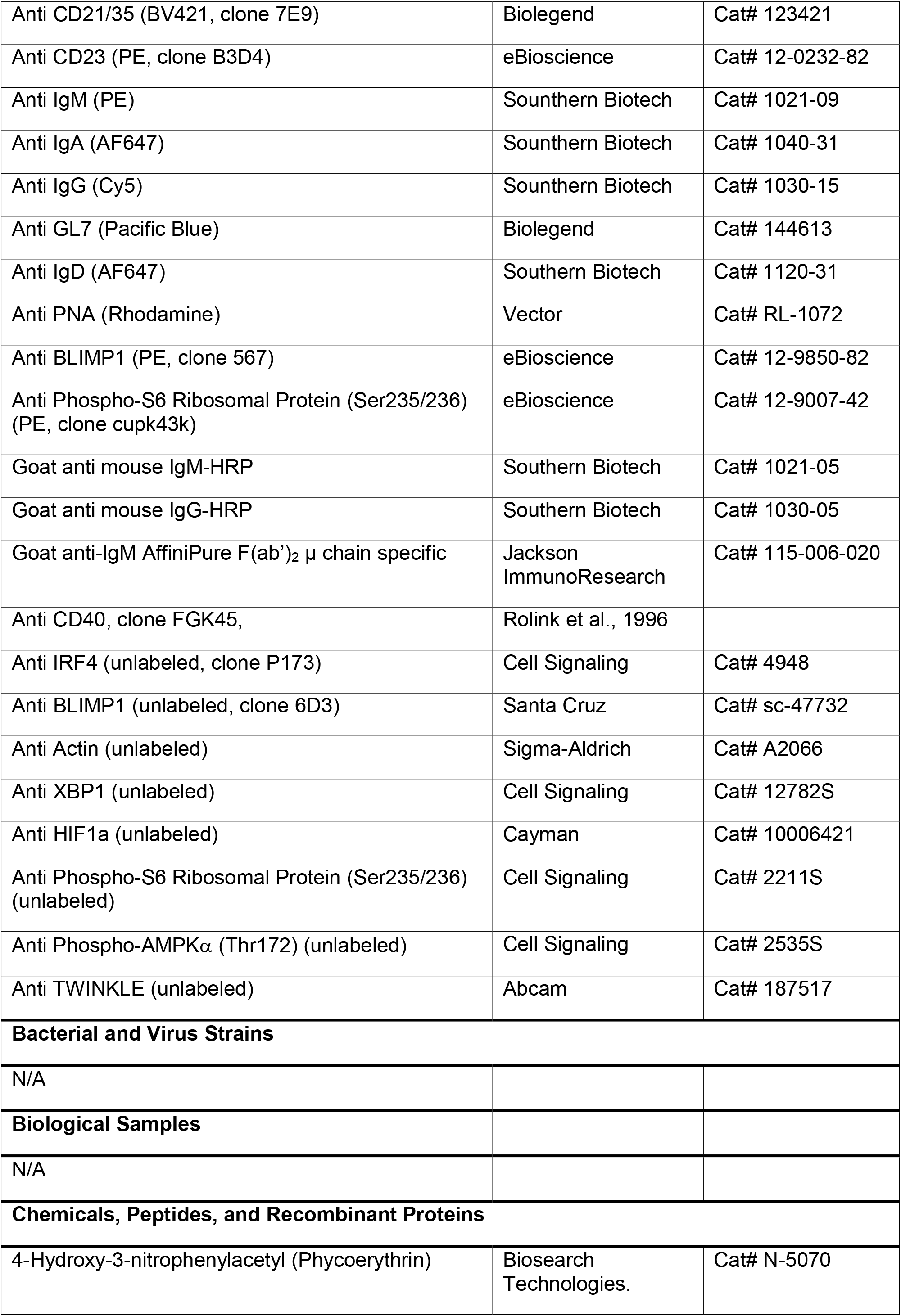
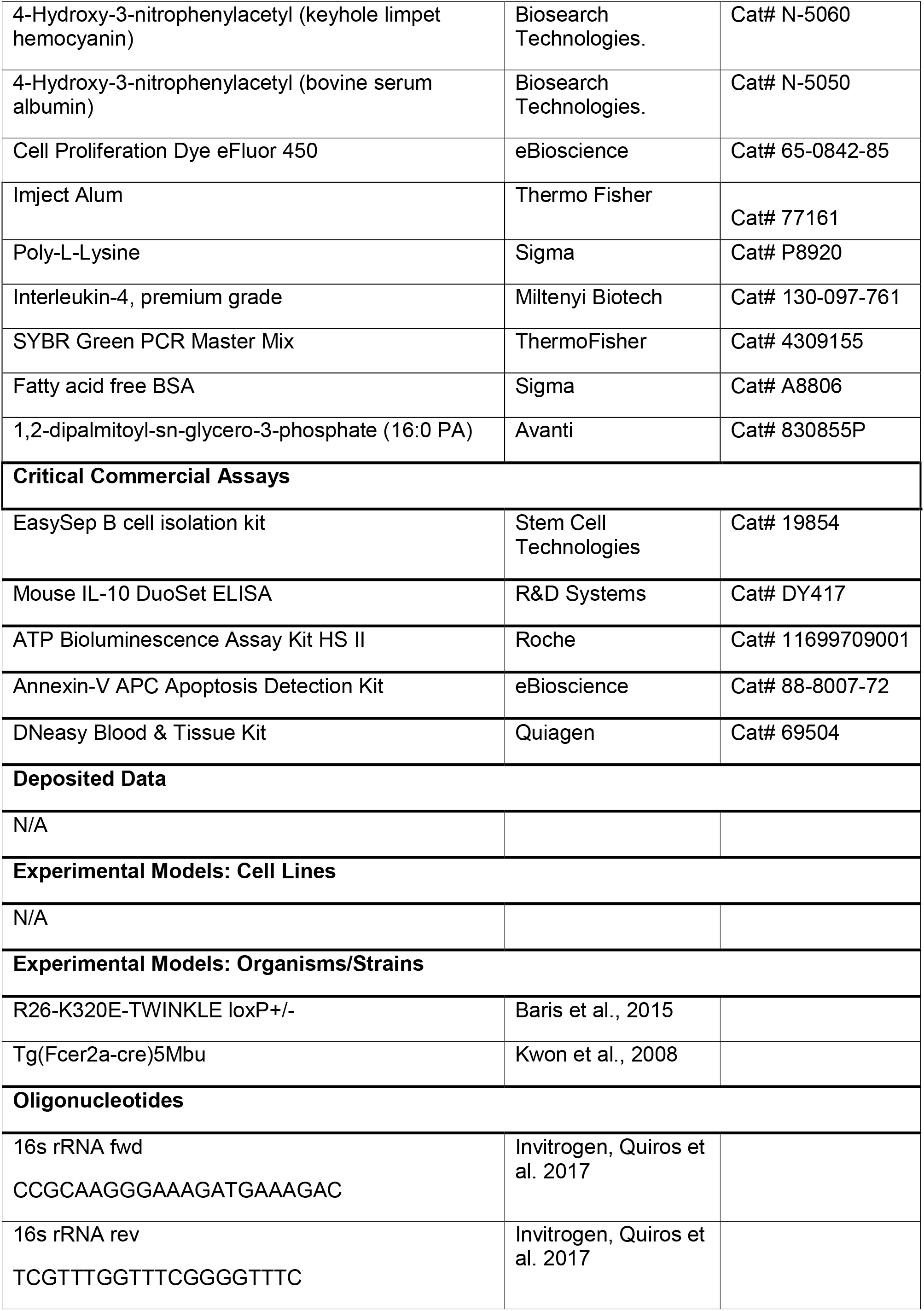
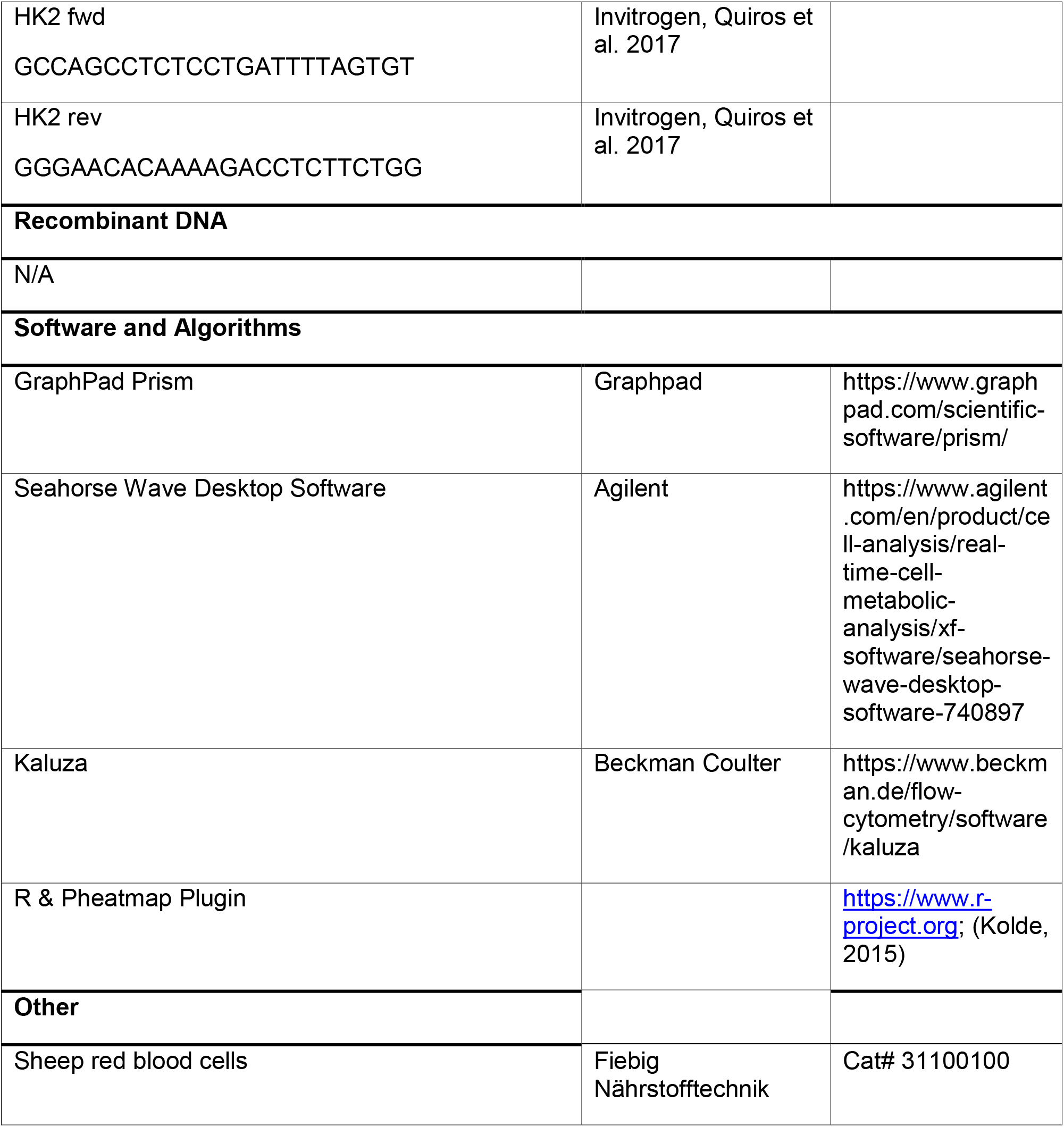

## STAR methods

### Resource Availability

#### Materials Availability

All unique reagents generated in this study are available from the Lead Contact.

#### Data and Code Availability

This study did not generate new resource datasets or code.

### Experimental Model and Subject Details

#### Animal models

All experimental procedures were done in agreement with animal protocols approved by the Government of Lower Franconia, Bavaria, Germany. Both female and male mice were used in the experiments. Mice were maintained on a 12-h light/dark cycle with free access to food and water according to governmental rules. All mice were in C57BL/6 backgrounds and between 8-12 weeks old. Sex-matched littermates or age and sex matched animals were used as controls. Mice were kept under pathogen free conditions in the mouse facility of the Nicolaus-Fiebiger-Center (Erlangen, Germany).

### Materials & Methods

#### Glycerophospholipid analysis

Glycerophospholipids (PC, PE, PI, PS, PG, PA) in B cells were analyzed by Nano-Electrospray Ionization Tandem Mass Spectrometry (Nano-ESI-MS/MS) with direct infusion of the lipid extract (*Shotgun Lipidomics*): 14 to 45 × 10^6^ cells were homogenized in 300 µl of Milli-Q water using the Precellys 24 Homogenisator (Peqlab, Erlangen, Germany) at 6.500 rpm for 30 sec. The protein content of the homogenate was routinely determined using bicinchoninic acid.

To 100 µl of the homogenate 400 µl of Milli-Q water, 1.875 ml of methanol/chloroform 2:1 (v/v) and internal standards (125 pmol PC 17:0-20:4, 132 pmol PE 17:0-20:4, 118 pmol PI 17:0-20:4, 131 pmol PS 17:0-20:4, 62 pmol PG 17:0/20:4, 75 pmol PA 17:0/20:4 Avanti Polar Lipids) were added. Lipid extraction and Nano-ESI-MS/MS analysis were performed as previously described (Kumar et al., 2015). Endogenous glycerophospolipids were quantified by referring their peak areas to those of the internal standards. The calculated glycerophospolipid amounts were normalized to the protein content of the tissue homogenate.

#### Mice

K320E-TWINKLE floxed mice (Baris et al., 2015) were crossed to CD23 CRE mice (Kwon et al., 2008) (kindly provided by Meinrad Busslinger) to generate DNT animals. DNT mice used in these experiments had the genetic background DNT^+/-^ CRE^+/-^ and CRE control mice were DNT^-/-^ CRE^+/-^. The WT animals used in this study were DNT^-/-^ CRE^-/-^ littermates. All mice were on the C57Bl/6 background.

#### T-dependent immunizations

Mice were injected with 100µg NP_29_-KLH (Biosearch Technologies) in Alum (ThermoScientific) in a 1:2 ratio (200µl total volume or in PBS after 6 weeks for boost immunization (*i.p*.).

#### Isolation of primary murine cells from spleen and bone marrow

Spleen was transferred in cold 2% FCS (in PBS) and gently passed through a 70µm cell strainer (BD) using the plunger of a 5ml syringe (BD). Femur and tibia were flushed with cold 2% FCS using a 27G cannula (BD). Cell suspensions were pelleted by centrifugation at 300xg for 5min at 4°C. Erythrocytes were lysed upon resuspension in red blood cell-lysis buffer (150mM NH4Cl, 10mM KHCO_3_, 100µM EDTA) for 5min at room temperature. The reaction was stopped by adding cold 2% FCS before centrifugation at 300xg for 5min at 4°C. The final cell suspensions were kept in cold 2% FCS after filtration through 30µm mesh filter (Sysmex).

#### Detection of surface antigens by flow cytometry

2×10^6^-4×10^6^ cells were pelleted in FACS tubes (Micronic) at 300xg for 5min at 4°C and resuspended in 50µl of unlabeled anti-CD16/32 Ab (10μg/ml in FACS-buffer (PBS, 2%FCS, 0.05% sodium azide)) for 15min on ice. Cells were washed once with FACS-buffer by centrifugation at 300xg for 5min at 4°C, resuspended in 50µl FACS-buffer containing the respective fluorochrome-coupled Abs and incubated for 20min on ice in the dark. Cells were washed with FACS-buffer by centrifugation at 300xg for 5min at 4°C. Data were acquired using a Gallios flow cytometer (Beckman Coulter). Analyses were performed using Kaluza version 1.3 and 2.1 (Beckman Coulter). Abs and other reagents are described in the key resources table.

#### Enzyme-linked immunosorbent assay (ELISA)

Serum samples from NP-KLH immunized mice (see above) were analyzed in duplicates serially diluted on 96-well flat-bottom microtiter plates (Greiner bio-one) coated with 1µg/ml NP4-BSA or NP_20_-BSA conjugates (Biosearch Technologies) in 50µl/well coating buffer (15mM Na_2_CO_3_, 35mM NaHCO_3_) overnight at 4°C. Captured NP-specific Abs were detected with goat anti-mouse IgM and IgG specific horseradish peroxidase (HRP) - coupled Abs (1:1000, Southern Biotech) and the ELISA was developed using TMB substrate reagent (BD OptEIA) and acid stop (0.5M H_2_SO_4_). Optical density (oD) was measured at 450nm on a FLUOstar Omega Microplate Reader (BMG Labtech) Plates were normalized using ELISA IgM and IgG standards as internal reference.

For analysis of serum from unimmunized mice and LPS culture supernatant, microtiter plates were coated with 1μg/ml goat anti-IgM/anti-IgG in coating buffer overnight at 4°C and then blocked for 1h at room temperature with 275μl/well of PBS, 2%FCS. Samples were analyzed in duplicates diluted serially 2-fold in PBS, 2%FCS for 1h at room temperature. Plates were washed 3 times with PBS, 0.05% Tween20 and then incubated with goat anti-mouse IgM-HRP/IgG-HRP in PBS, 2%FCS. After washing, plates were developed using developed using TMB substrate reagent (BD OptEIA) and acid stop (0.5M H_2_SO_4_). Optical density (oD) was measured at 450nm on a FLUOstar Omega Microplate Reader (BMG Labtech). Plates were normalized using ELISA IgM and IgG standards as internal reference.

ELISA for detection of IL10 in culture supernatant was performed using the Mouse IL-10 DuoSet ELISA (R&D Systems) according to manufacturer’s instructions.

#### Western Blot

Proteins were electrophoresed on 10% SDS polyacrylamide gels and transferred to a nitrocellulose membrane (45 min, 400 mA). Transfer efficiency was determined by Ponceau S. Membranes were blocked in 5% skimmed milk powder in Tris-buffered saline (TBS) containing 0.1% Tween-20 (TBST) and probed with the respective antibody at 4°C overnight diluted in 3% BSA in PBS, containing 0.1% Tween-20 and 0.1% Sodium Azide. Membranes were washed in TBST and incubated with anti-mouse/rabbit/rat IgG HRP conjugate diluted in 5% skimmed milk powder in TBST for 1 hour at room temperature. After washing, blots were developed by enhanced chemiluminescence. Quantification of western blot bands was performed by densitometry analysis with ImageJ. Therefore, scanned X-ray films were converted to 8-bit type and color inverted. Band intensities were measured, and background of X-ray films was subtracted. Protein expression was normed to β-Actin as loading control.

#### Purification of murine B lymphocytes from spleen

B cells were enriched from splenic cell suspensions using the EasySep Mouse B cell isolation negative selection kit (EasySep #19854, Stemcell Technologies) according to the manufacturer’s instructions. In short, spleen cells were resuspended in MACS buffer (PBS, 2% FCS, 2mM EDTA), surface blocked with rat serum and immuno-magnetically enriched for B cells. Purity of isolated B cells was verified by surface stain for CD19. Usually, an enrichment of > 95% was achieved.

#### *In vitro* cultivation of primary murine B cells

Splenic B cells were cultured with a starting concentration of 0,5×10^6^ cells/ ml in R10 medium (RPMI1640, 10% fetal calf serum (FCS), 2mM glutamate, 1mM sodium pyruvate, 50 U/ml penicillin G, 50μg/ml streptomycin, 50μM β-mercaptoethanol) for 72h at 37°C and 5% CO_2_, supplemented with 10µg/ml LPS. For *in vitro* class switch recombination cells were seeded at 0,1×10^6^ cells/ ml in R10 medium for 96h, supplemented with 5ng/ml TGF beta, 5nM retinoic acid, 10µg/ml α-CD40, 10µg/ml LPS, 100U/ml IL4 and 10ng/ml IL5.

#### Extracellular Flux Analysis

The day prior to the assay cell plates were coated with 10µg/ml Poly-L-Lysine in 1x TE buffer. Day 3 LPS blasts were seeded at a density of 2,5×10^5^ cells/well, measured at least in triplicates. Mito Stress Test was performed according to manufacturer’s instructions.

#### Intracellular ATP measurement

Level of intracellular ATP of 5×10^4^ cells per sample was determined using the ATP Bioluminescence Assay Kit HS II (Roche) according to manufacturer’s instructions.

#### DNA extraction and RT-qPCR

Cell pellets were resuspended in PBS and DNA was isolated using the DNeasy Blood & Tissue Kit (Quiagen), according to manufacturer’s instructions. qPCR was performed using SYBR Green I-dTTP (Eurogentec) using the Applied Biosystems 7500 real-time PCR system. Samples were analyzed in triplicates and 16s rRNA copies, representing mtDNA copies, were normalized to the HK2 gene.

#### Liposome production

Liposome production was based on a recently published method (S. T. Poschenrieder, 2016) originally developed for the production of polymersomes, which are the artificial counterparts of liposomes. Here, 11.4 mL of vesicle buffer (10 mM TRIS-HCl, 150 mM NaCl, pH 8.0) were poured into unbaffled miniaturized stirred tank reactors (BioREACTOR48, 2mag, Munich, Germany) and stirred at 4,000 rpm at 25°C using an S-type stirrer. Then, 600 µL of the dissolved phospholipid 16:0 PA (1,2-dipalmitoyl-sn-glyero-3-phosphate) (10 mM in chloroform) were injected under stirring, leading to a whitish highly dispersed emulsion. The reactor shell, originally made of polystyrene, has been replaced by borosilicate glass, in order to avoid damaging by the chloroform. The process was continued, until the solution became transparent, indicating the evaporation of the chloroform, which was the case after 6 hours. Subsequently, the solution was centrifuged at 13,000 rpm in a tabletop centrifuge to remove precipitates. The liposomes in the supernatant were then concentrated to 1,5 mg mL^-1^ by centrifugation for 50 min at 50,000 g and resuspending of the resulting pellet in the appropriate amount of vesicle buffer. The quality of the liposomes during and after the production process was monitored via dynamic light scattering using a Zetasizer NS (Malvern, Worcestershire, UK) as reported (S. T. Poschenrieder, 2016).

#### Lactate and Glucose measurement of culture supernatant

Supernatant of LPS activated B cells was harvested after 3 days and analyzed with the SUPER GL compact analyzer according to the manufacturer’s instructions.

#### Immunohistology

Mice were immunized with SRBCs, sacrificed after 10 days and spleens were embedded in OCT medium (Tissue-Tek). Cryotome sections of 8µm were prepared, fixed in acetone, blocked (with 20 µg/ml α-CD16/23 in 1% PBS, 10% FCS, 2% BSA) and stained with α-IgD-AF647, PNA-Rhodamine and α-GL7-PacificBlue.

#### Metabolomics

Splenic B cells were isolated, activated with LPS and viable cells, only GFP^+^ for DNT, were sorted after 3 days using flow cytometry. Perchloric acid extraction was performed and metabolic profiles as previously published (Hofmann et al., 2011).

#### Electron microscopy

Viable cells were collected by ficoll density gradient centrifugation and were fixed overnight in 2.5% glutaraldehyde (Carl Roth, 4995.1) in PBS. Cells were further processed and analyzed, as described previously (Steinmetz et al., 2020).

#### Quantification and statistical analysis

Values were assessed for Gaussian distribution using the Shapiro-Wilk normality test. Mann-Whitney test was used for non-Gaussian distributed data sets. Datasets revealing Gaussian-like distribution were assessed by Student’s t test. Differences between the analyzed groups were considered to be statistically significant with p-values <0.05. *, p<0.05, **, p < 0.01, ***, p < 0.002, ****, p < 0.0004. Data were analyzed using Prism (GraphPad) and R using the Pheatmap plugin (Kolde, 2015).

---

**Figure S1.**
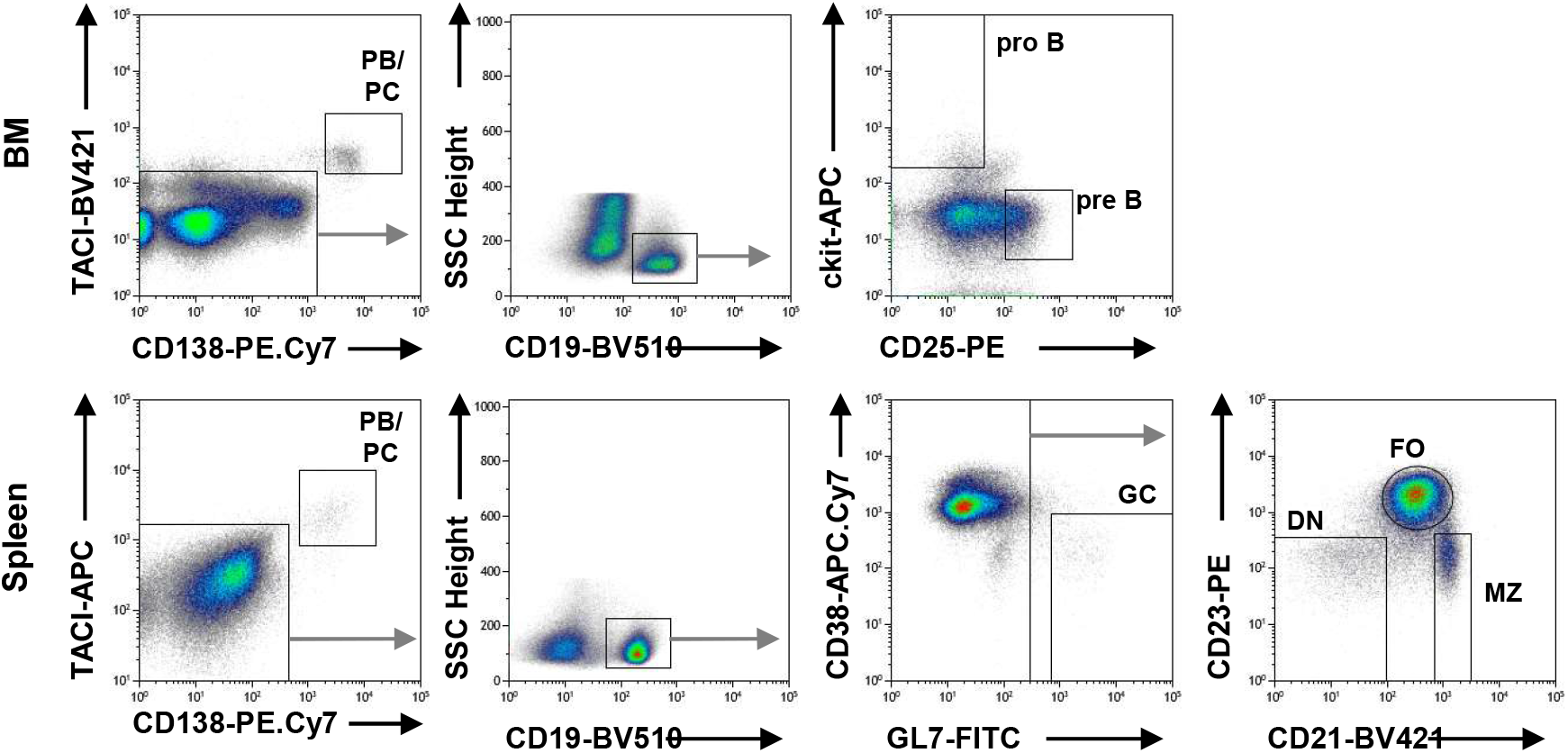
Flow cytometric cell sorting of murine B cell subsets. WT mice were immunized with SRBCs and B cell subsets were sorted using flow cytometry from the bone marrow and spleen after 10 days. Bone marrow (top): PCs are defined as TACI^+^CD138^+^, pro B cells are CD19^+^ckit^+^ and pre B cells are CD19^+^CD25^+^. Spleen (bottom): PCs are TACI^+^CD138^+^, GCs are CD19^+^CD38^low^GL7^high^, FOs are CD19^+^CD21^+^CD23^+^, MZs are CD19^+^CD21^+^CD23^low^ and DN are CD19^+^CD21^-^CD23^-^. Related to Figure 1.

**Figure S2.**
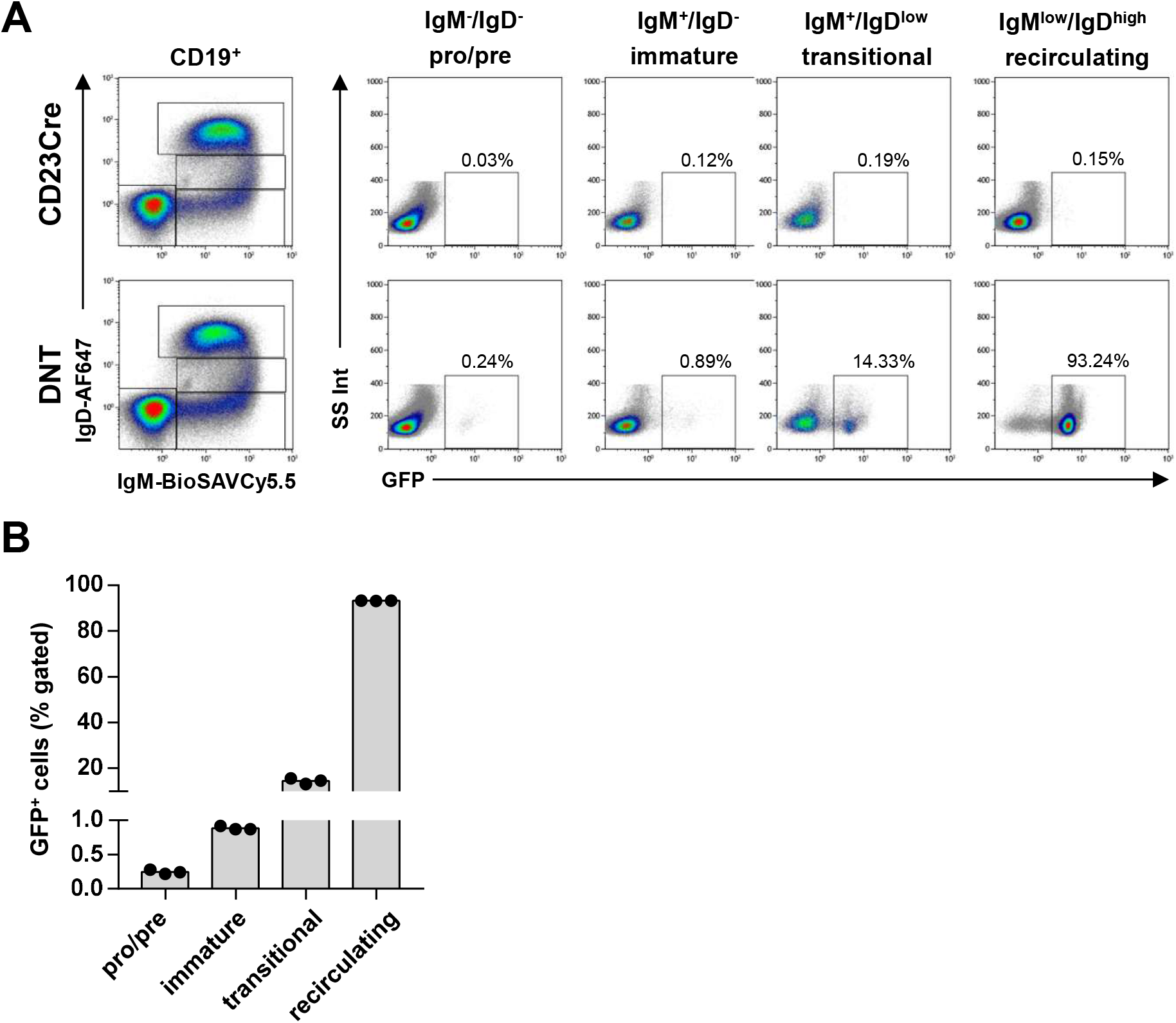
Onset of GFP expression during B cell development in DNT mice. A, Bone marrow cells of CD23CRE and DNT mice pregated on CD19^+^ were stained with anti IgM and anti IgD antibodies. GFP positive cells of the indicated B cell populations are depicted as dot blots. B, Frequencies of GFP positive cells in the indicated populations defined in A. Each dot represents one mouse. Related to Figure 1.

**Figure S3.**
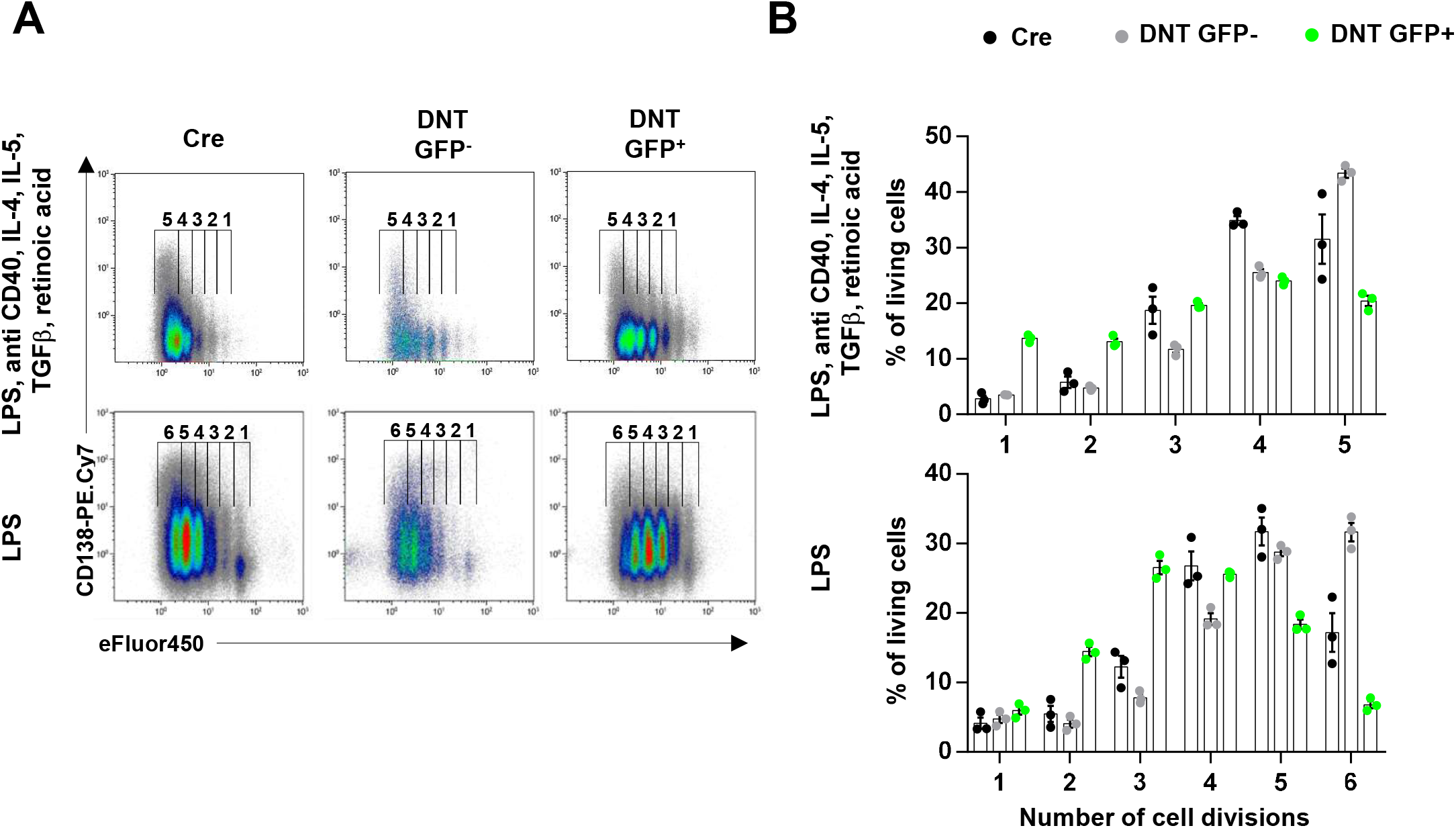
Impaired proliferation of T-dependently and T-independently activated DNT B cells. A, Splenic B cells from CD23CRE and DNT mice were labelled with eFluor450, stimulated with LPS, anti CD40, IL-4, IL-5, TGFb, retinoic acid for 4 days or with LPS alone for 3 days, stained with anti CD138 antibody and analyzed by flow cytometry. B, CD138 expressing B cells relative to the number of cell divisions of CD23CRE or DNT mice is depicted. Black symbols represent CD23CRE B cells, green symbols show GFP^+^ B cells of DNT mice and grey symbols show GFP^-^ B cells of DNT mice from the same culture. Related to Figure 5.

**Figure S4.**
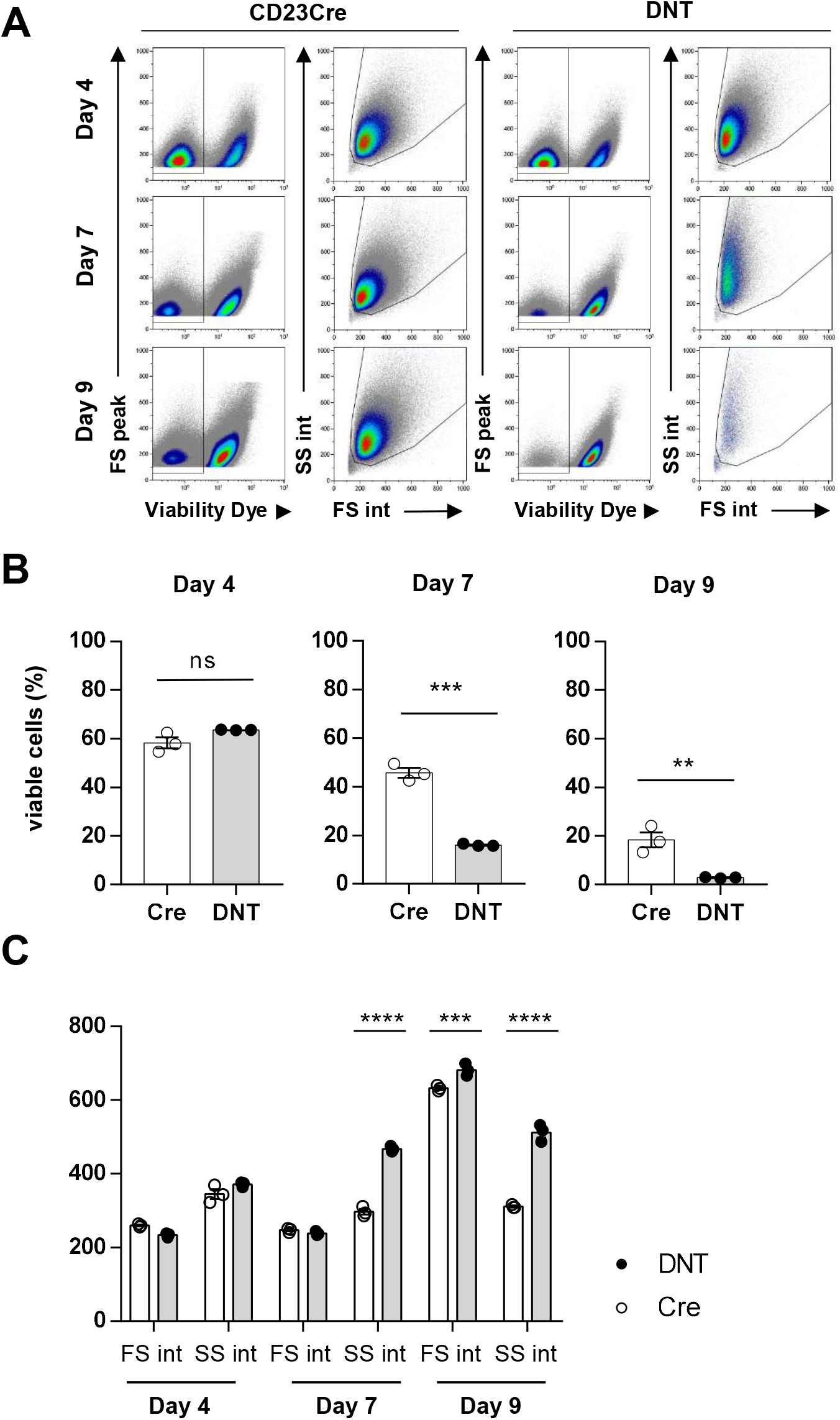
Impaired survival and morphological changes activated DNT B cells. A, Splenic B cells from CD23CRE and DNT mice were stimulated with LPS, anti CD40, IL-4, IL-5, TGFβ, retinoic acid for 4, 7 and 9 days, stained with a viability dye and analyzed by flow cytometry. Merged dot plots of 3 mice per genotype are shown. B, Quantification of viable cells according to the gating in A. C, Quantification of the forward and side scatters of viable cells shown in A. Related to Figure 5.

**Figure S5.**
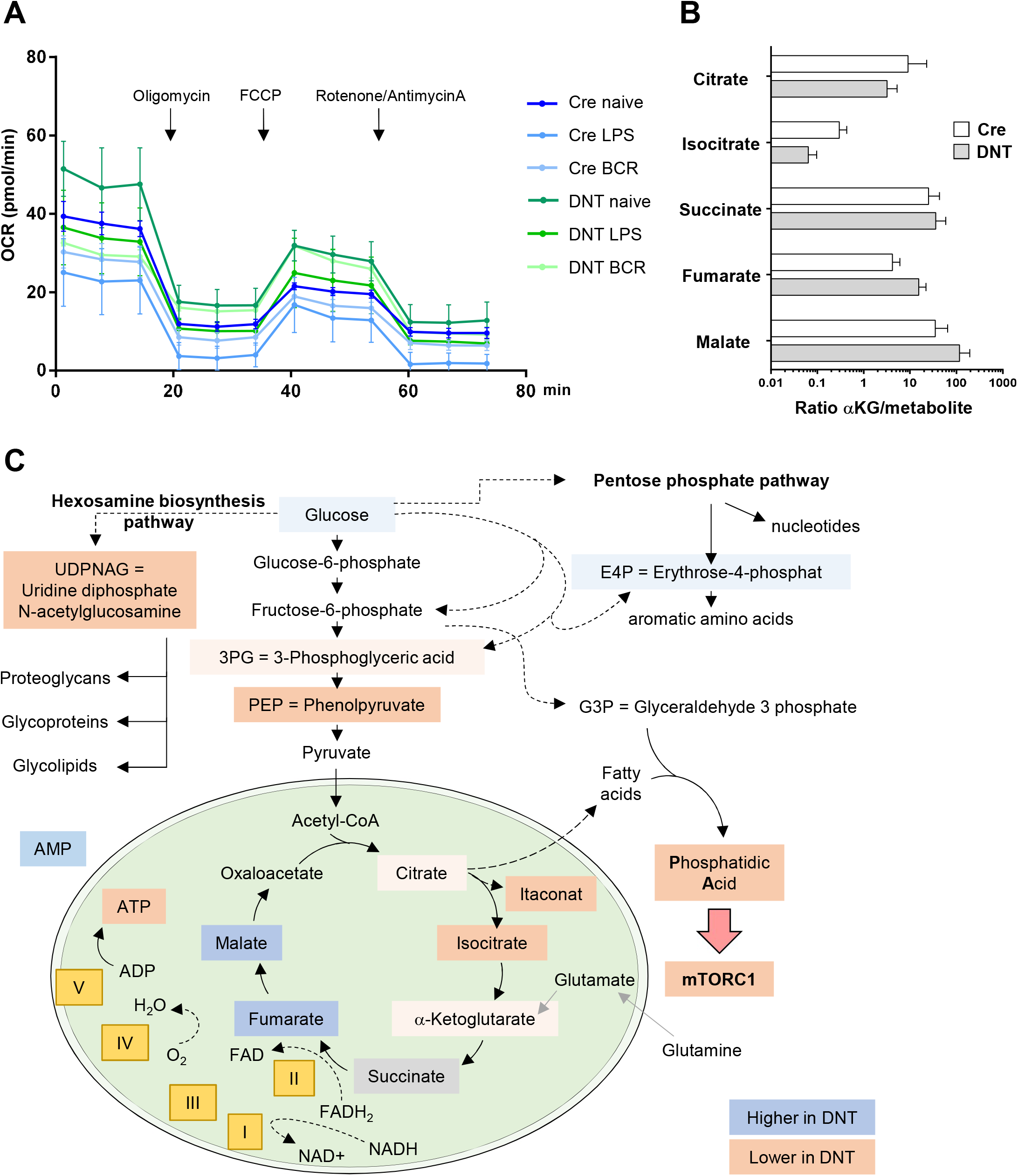
Metabolic alterations of resting and LPS activated DNT B cells. A, Splenic B cells from CD23CRE and DNT mice were left non-activated, stimulated with LPS or with anti B cell receptor antibody for 6h and extracellular flux was analyzed. The oxygen consumption rate (OCR) was measured basally and after injection of oligomycin, FCCP and rotenone + antimycin A. Symbols represent means of 2-4 mice, error bars are ± SEM. B, Ratio of α-Ketoglutarate to upstream and downstream metabolites of the TCA cycle in 3d LPS stimulated CRE and DNT B cells. C, Schematic of glycolysis, the hexosamino biosynthesis and pentose phosphate pathways and the TCA cycle. Roman letters indicate OxPhos complexes I-V. Metabolites decreased in DNT B cells are shown in orange colour, increased metabolites are indicated in blue colour, and unchanged in grey. G3P, Glyceraldehyde-3-phosphate, 3PG, 3-Phosphoglycerate, E4P, Erythrose-4-Phosphate, NAD, Nicotinamide adenine dinucleotide, NADPH, Nicotinamide Adenine Dinucleotide phosphate, FAD, Flavine Adenine Dinucleotide, FADH_2_, Flavine Adenine Dinucleotide Dihydrate. Related to Figure 7.

**Figure S6.**
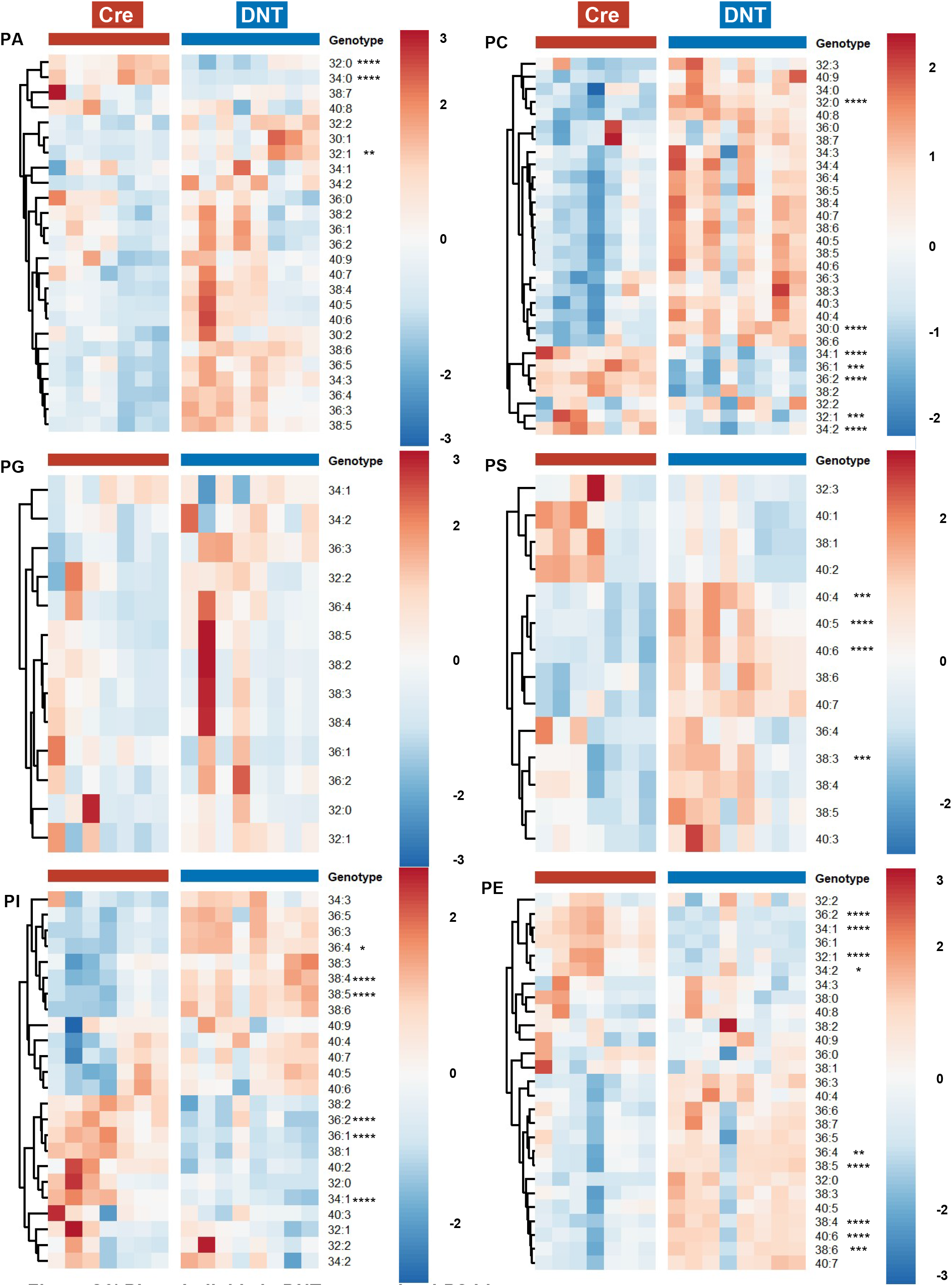
Phospholipids in DNT expressing LPS blasts. Splenic B cells from CD23CRE and DNT mice were stimulated with LPS for 3d. Glycerophospholipids of FACS sorted B cells (Gating: DNT: viable and GFP^+^, CD23CRE: viable) were analysed via direct infusion MS/MS (*Shotgun Lipidomics*). The relative abundance (Mol%) of subspecies of phosphatidic acid (PA), phosphatidylglycerol (PG), phosphatidylinositol (PI), phosphatidylcholine (PC), phosphatidylethanolamine (PE) and phosphatidylserine (PS) is depicted as heatmap. Each symbol represents one mouse. The first number depicts the total number of carbon atoms, the second number the total number of double bonds within the two fatty acyl chains. Combined results from 2 experiments, N=2, n=3-5; statistics calculated using 2-way ANOVA. Related to Figure 8.

**Figure S7.**
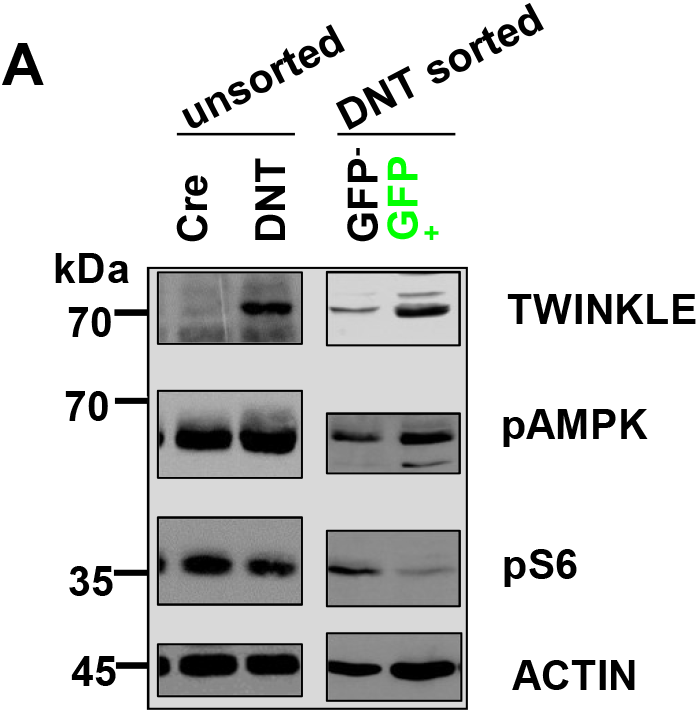
pAMPK and pRPS6 expression in DNT expressing B cells. Unsorted splenic B cells from CD23CRE and DNT mice were stimulated with LPS for 3d. Cell lysates were separated by 10% SDS-PAGE, transferred to nitrocellulose and stained with antibodies as indicated on the right (left). Right: Splenic B cells from DNT mice were stimulated with LPS for 3d. GFP^+^ and GFP^-^ cells were sorted out. Cell lysates were separated by 10% SDS-PAGE, transferred to nitrocellulose and stained with antibodies as indicated on the right. Molecular mass standards are shown on the left (kDa). Related to Figure 8.

**Figure S8.**
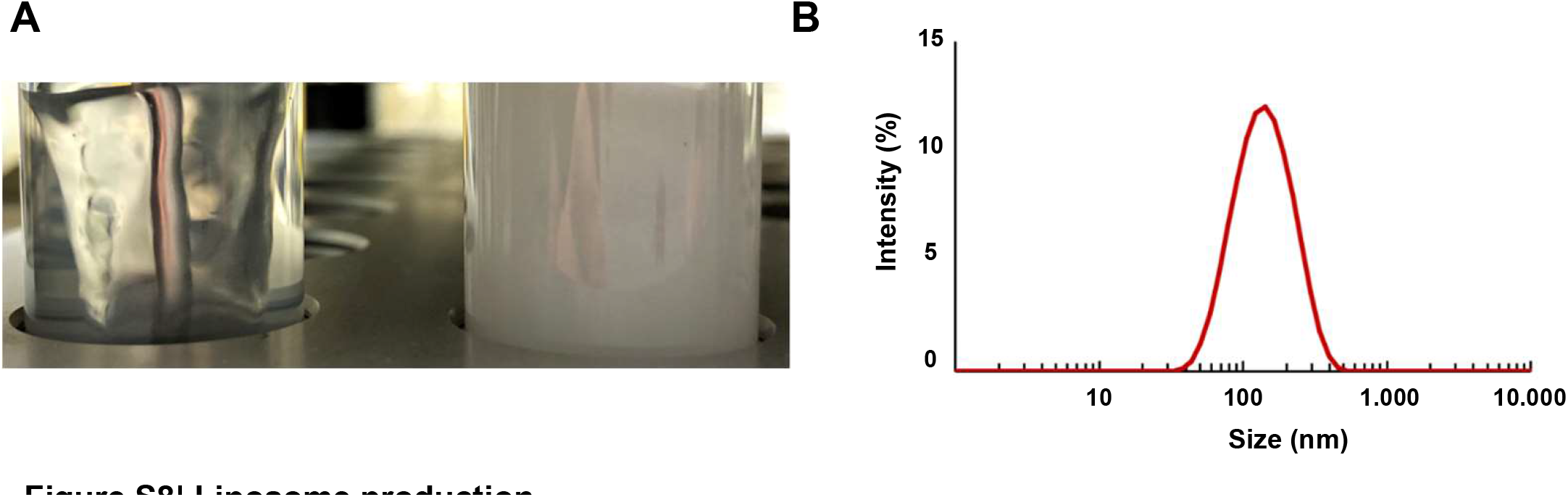
Liposome production. A, Liposome production in miniaturized stirred tank reactors. On the left: Liposomes in aqueous buffer after evaporation of the chloroform (6 hours after injection). On the right: Dispersed whitish emulsion directly after injection of the phospholipid 16:0 PA, dissolved in chloroform. B, Intensity based particle size distribution of the liposomes after a final process time of 6 hours, shown exemplarily for one of the liposome samples used in this study. The sample showed a poly dispersity index (PDI) of 0.167 and a z average cumulants mean of 122 nm. Related to Figure 9.

